# Viability of AMURA biomarkers from single-shell diffusion MRI in Migraine Clinical Studies

**DOI:** 10.1101/2022.04.01.486661

**Authors:** Carmen Martín-Martín, Álvaro Planchuelo-Gómez, Ángel L. Guerrero, David García-Azorín, Antonio Tristán-Vega, Rodrigo de Luis García, Santiago Aja-Fernández

## Abstract

Diffusion Tensor Imaging (DTI) is the most employed method to assess white matter properties using quantitative parameters derived from diffusion MRI, but it presents known limitations that restricts the evaluation of complex structures. The objective of this study was the assessment of complementary diffusion measures extracted with a novel approach, Apparent Measures Using Reduced Acquisitions (AMURA), in comparison with DTI in clinical studies. Fifty healthy controls, 51 episodic migraine and 56 chronic migraine patients underwent single-shell diffusion MRI. Three DTI-based and eight AMURA-based parameters were compared between groups with tract-based spatial statistics. On the other hand, following a region-based analysis, the measures were assessed for multiple subsamples with diverse reduced sample sizes and their stability was evaluated with the coefficient of quartile variation. Compared to DTI, statistically significant differences in additional regions and group comparisons were found with AMURA-based measures. Additionally, some AMURA-based measures showed a remarkable discriminative power for quite reduced sample size and, simultaneously, lower stability (higher coefficient of quartile variation) than DTI-based measures. These findings suggest that AMURA presents favorable characteristics to identify differences of specific microstructural properties between clinical groups in regions with complex fiber architecture and lower dependency on the sample size or assessing technique than DTI.

## 1. INTRODUCTION

Diffusion Magnetic Resonance Imaging (dMRI) is an imaging modality employed to assess diverse *in vivo* physiological and pathological conditions of the human body in clinical studies. It has been widely used in the study of the brain and neurological disorders ^1–4^. It allows the characterization of the diffusivity of water molecules within the tissue, providing information about the microscopic configuration and structural connectivity of the brain, especially inside the white matter (WM).

The most relevant feature of dMRI is its ability to measure directional variance, i.e., anisotropy, which, inside the brain, is related to connectivity between areas. The most common methodology to estimate the anisotropy is via the diffusion tensor (DT) ^5,6^.

In order to carry out clinical studies, the information provided by the DT must be translated into some scalar measures that describe different features of diffusion within every voxel. That way, metrics like fractional anisotropy (FA) were defined and widely employed to characterize damaged tissues in multiple neurological and psychiatric disorders ^7–10^. However, from the early stages of DT imaging (DTI), it was clear that the Gaussian assumption oversimplifies the diffusion process.

In the past few decades, many techniques were proposed to overcome the limitations of DTI, including the acquisition of larger amounts of diffusion data ^11,12^. Most of these techniques rely on the estimation of more advanced diffusion descriptors, such as the Ensemble Average diffusion Propagator (EAP), which is the probability density function of the motion of the water molecules within a voxel ^13–15^.

A complete analysis of the EAP requires many diffusion-weighted images (DWI) with several (moderate to high) b-values in a multi-shell acquisition. The information provided by the EAP is usually adapted to scalar measures that describe different aspects of diffusion. The most frequently employed measures are the return-to-origin probabilities (RTOP), return-to-plane probabilities (RTPP), return-to-axis probabilities (RTAP) and the propagator anisotropy (PA) ^13,16– 19^.

The accurate estimation of these measures requires the calculation of the EAP, which commonly involves: (1) long acquisition times; (2) several shells with large b-values, which may be difficult to acquire in many commercial MRI scanners; and (3) heavy computational burdens with very long processing times. These three issues have hindered the general adoption of EAP-related metrics in the clinical routine, despite the growing interest in the exploration of their potential applicability ^20–23^.

To overcome these limitations and facilitate the widespread use of advanced diffusion metrics in clinical studies, a new approach called *Apparent Measures Using Reduced Acquisitions* (AMURA) has been recently proposed ^24–26^. The method allows the estimation of diffusion measures such as RTOP, RTAP and PA, while reducing the number of necessary samples and the computational cost. AMURA can mimic the sensitivity of EAP-based measures to microstructural changes when a reduced amount of data distributed in a few shells (even one) is available. To do so, AMURA assumes a prior model for the behavior of the radial q–space instead of trying to numerically describe it, yielding simplified expressions that can be computed easily even from single-shell acquisitions. However, since the assumed mono-exponential model only holds within a limited range around the measured b-value, the measures derived this way must be seen as *apparent values* at a given b-value, related to the original ones but dependent on the selected shell.

One additional advantage of AMURA is that it can be easily integrated into the processing pipeline of current existing single-shell dMRI protocols and databases to unveil anatomical details that may remain hidden in traditional DT-based studies. AMURA has proved its potential in some exploratory studies with clinical data focusing on Parkinson and Mild Cognitive Impairment ^24,26^, as well a recent clinical study on migraine ^27^.

In this work we aim to assess the viability of different diffusion descriptors extracted with AMURA for the study of a neurological disorder in DTI-type datasets. Note that, initially, AMURA was designed to work with b-values over 2000 s/mm^2^, since the effects measured with RTOP, RTPP and RTAP were better showed at higher values of b. However, results in clinical data have shown its potential at lower b-values ^25^. Thus, we will explore the viability of these technique to model DTI-type acquisitions, i.e., dMRI datasets acquired with those protocols usually employed for the estimation of DTI and its derived parameters, such as FA or MD. These acquisitions are commonly single-shell, and only include one non-zero b-value, usually in the order of b=1000 s/mm^2^.

We have selected migraine as a case study. Migraine is an attractive pathology for the evaluation of the quality of alternative diffusion metrics, since the differences between patients and controls that have been found using dMRI in the literature are scarce and subtle ^28^. In migraine, differences are usually hard to find in comparison with other disorders such as schizophrenia or Alzheimer’s disease, and they require a large number of subjects per group and good quality data. Thus, migraine will allow us to check the capability of different techniques to detect subtle changes.

Migraine is a disabling primary disorder characterized by recurrent episodes of headache, which usually last 4-72 hours and present at least two of the following four characteristics: moderate to severe pain intensity, unilateral location, pulsating quality, and aggravation with physical activity (Third edition of the International Classification of Headache Disorders, ICHD-3). A common distinction when studying migraine is made between episodic migraine (EM), in which patients suffer from headache less than 15 days per month, and chronic migraine (CM), in which patients suffer from headache at least 15 days per month.

A recent study identified statistically significant differences in migraine using advanced diffusion measures calculated with AMURA ^27^. This study identified higher RTOP values in CM patients compared to EM, and lower RTPP values in EM compared to HC.

Given the fact that AMURA-derived measures have shown promising results for the characterization of subtle WM changes in migraine, the main objective of this study was the assessment of the reliability and the robustness of AMURA metrics acquired with a typical acquisition employed in a clinical context. Our purpose is to validate the viability of these metrics for clinical studies even when acquisition protocols are suboptimal for this methodology. Specifically, we will use migraine as a case study and DTI-type acquisitions, where only one shell is acquired at b=1000 s/mm^2^.

## 2. MATERIALS AND METHODS

### 2.1. Advanced diffusion measures from single shell acquisitions: AMURA

AMURA was proposed ^24^ as a methodology to calculate advanced diffusion metrics from reduced acquisitions compatible with commercial scanners and general clinical routine. It allows the estimation of different diffusion-related scalars using a lower number of samples with a single-shell acquisition scheme. AMURA considers that, if the amount of data is reduced, a restricted diffusion model consistent with single-shell acquisitions must be assumed: the (multi-modal) apparent diffusivity does not depend on the b-value, so that a mono-exponential behavior is observed for every spatial direction. According to ^29^, in the mammalian brain, the mono-exponential model is predominant for values of b up to 2000 s/mm^2^ and it can be extended to higher values (up to 3000 s/mm^2^) if appropriate multi-compartment models of diffusion are employed. Appendix A shows a graphical abstract figure of the calculation of AMURA.

This methodology allows shorter MRI acquisitions and very fast calculation of scalars. Since the mono-exponential model only holds within a limited range around the measured b-value, the measures derived this way must be seen as apparent values at a given b-value, related to the original ones but dependent on the selected shell. The AMURA metrics used in this work are ^24– 26^:

1. Return-to-origin probability (RTOP), also known as probability of zero displacement, it is related to the probability density of water molecules that minimally diffuse within the diffusion time τ.
2. Return-to-plane probability (RTPP), which is a good indicator of restrictive barriers in the axial orientation.
3. Return-to-axis probability (RTAP), an indicator of restrictive barriers in the radial orientation.
4. Apparent Propagator Anisotropy (APA), an alternative anisotropy metric. It quantifies how much the propagator diverges from the closest isotropic one.
5. Diffusion Anisotropy (DiA), an alternative derivation of APA.
6. Generalized Moments, specifically we will consider the full moments of order 2 (q-space Mean Square Displacement, qMSD) and ½ (ϒ ^1/2^).

Expressions for the calculation of the different AMURA metrics can be found in Appendix B. Finally, note that, since the mono-exponential model assumed by AMURA only holds within a limited range around the measured b value, these metrics must be considered as *apparent values* at a given b-value.

### 2.2. Dataset

#### 2.2.1. Participants

The sample of this study was originally composed of 56 patients with CM, 54 patients with EM and 50 healthy controls (HC) that participated in previous studies ^28,30^. Three patients with EM were discarded due to misregistration errors.

Inclusion criteria included diagnosis of EM or CM following the ICHD-3 (all the available versions), stable clinical situation, and first screening related to migraine just before the recruitment. Exclusion criteria were use of preventive treatments before the MRI acquisition, migraine onset in people older than 50 years, recently developed migraine (less than one year), frequent painful conditions, psychiatric and neurological disorders different to migraine, and pregnancy. Further details are available at ^28^.

The local Ethics Committee of Hospital Clínico Universitario de Valladolid approved the study (PI: 14-197). Additionally, all participants read and signed a written consent form prior to their participation.

#### 2.2.2. MRI acquisition

For patients with migraine, the images were acquired at least 24 hours after the last migraine attack and before two weeks after the clinical visit to the headache unit. High resolution 3D T1-weighted followed by DWI were acquired using a Philips Achieva 3T MRI unit (Philips Healthcare, Best, The Netherlands) with a 32-channel head coil.

The acquisition of T1-weighted images was carried out using a Turbo Field Echo sequence with the following parameters: repetition time (TR) = 8.1 ms, echo time (TE) = 3.7 ms, flip angle = 8°, 256x 56 matrix size, spatial resolution of 1×1×1 mm^3^ and 160 sagittal slices covering the whole brain.

The acquisition parameters for DWI were TR = 9000 ms, TE = 86 ms, flip angle = 90°, 61 diffusion gradient orientations, one baseline volume, b-value = 1000 s/mm^2^, 128×128 matrix size, spatial resolution of 2×2×2 mm^3^ and 66 axial slices covering the whole brain.

All the images were acquired in the same session with a total acquisition time of 18 minutes.

### 2.3. Analysis of the data

#### 2.3.1. dMRI preprocessing

Image preprocessing steps consisted of 1) denoising based on the Marchenko-Pastur Principal Component Analysis procedure ^31^, 2) eddy currents and motion correction and 3) correction for B1 field inhomogeneity. The MRtrix software ^32^ was employed to carry out these steps, using the *dwidenoise, dwipreproc* and *dwibiascorrect* tools ^31,33–35^. Further, a whole brain mask for each subject was acquired with the *dwi2mask* tool ^36^.

#### 2.3.2. Diffusion measures estimation

Two groups of diffusion measures were extracted. The former group is composed of three DTI classical metrics: FA, MD, AD and RD. These measures were estimated at each voxel using the *dtifit* tool from the FSL software ^37^. FA measures the degree of anisotropy in the diffusion of water molecules inside each voxel, which reflects the degree of directionality of water diffusivity. MD is the average magnitude of water molecules diffusion. AD is the principal direction of WM fibers of the water diffusion. RD describes the perpendicular diffusion of the principal direction ^38^.

The latter group includes the seven proposed q-space metrics calculated with AMURA: RTOP, RTAP, RTPP, APA, qMSD, DiA and ϒ^1/2^. The measures were calculated using *dMRI-Lab* (available at www.lpi.tel.uva.es/dmrilab) and MATLAB 2020a. AMURA measures rely on the expansion of spherical functions at a given shell in the basis of spherical harmonics (SH). Even SH orders up to 6 were fitted with a Laplace-Beltrami penalty λ = 0.006. A visual comparison of the DTI and AMURA measures is shown in Figure 1.

**Figure 1.**
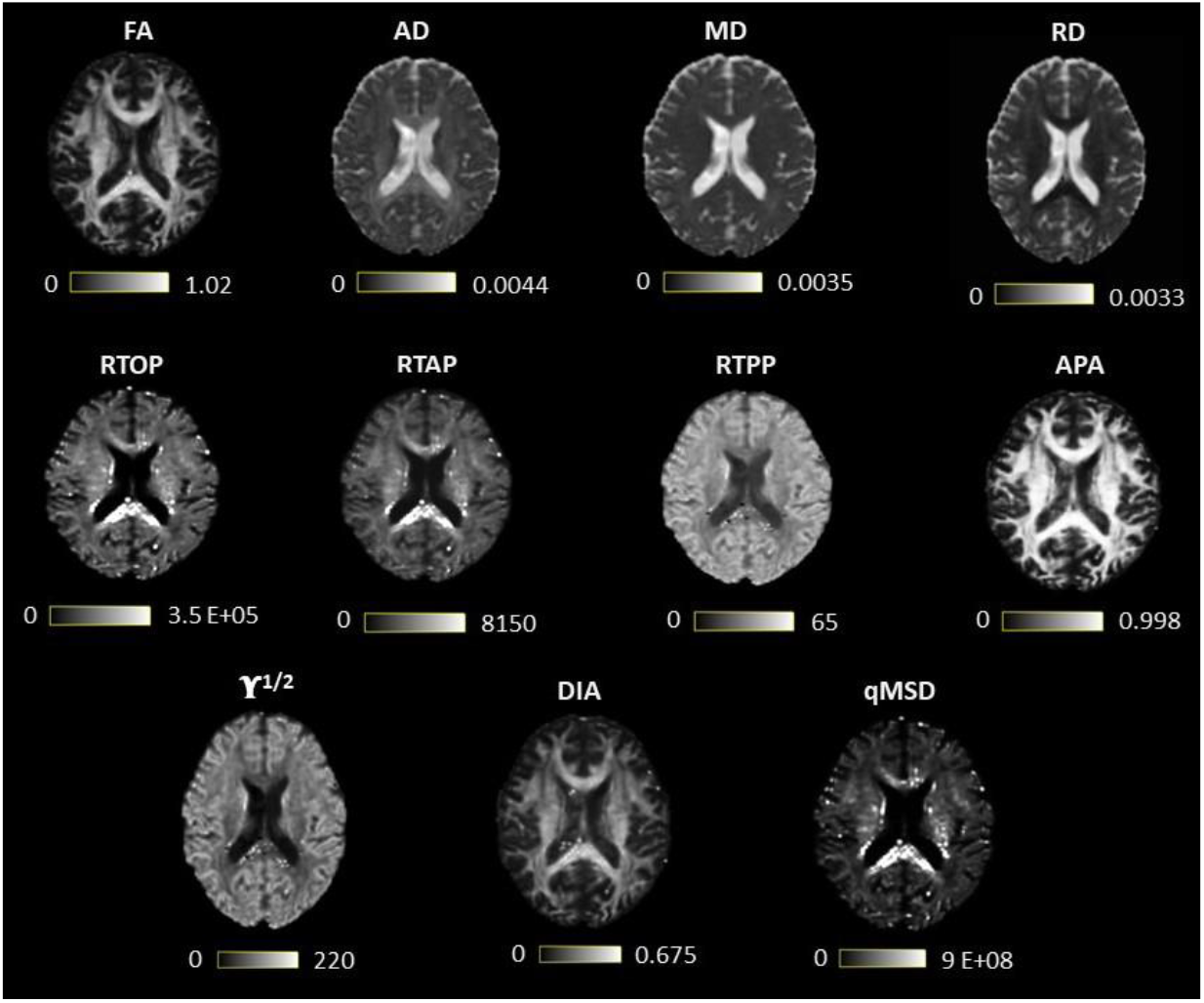
Visual comparison of diffusion tensor imaging (DTI) and measures from apparent measures using reduced acquisitions (AMURA). The first row contains the DTI measures, and the last two, the AMURA metrics.

### 2.4. Statistical analysis

#### 2.4.1. TBSS & statistical analysis

Tract-based spatial statistics (TBSS) was used to analyze diffusion measures differences in WM tracts between the three groups of interest: CM, EM and HC^38^. Forty-eight WM tracts were delimited according to the Johns Hopkins University ICBM-DTI-81 White Matter Atlas (JHU WM) ^39,40^. TBSS steps included nonlinear registration to the Montreal Neurological Institute (MNI) space, generation of a white matter skeleton from the mean FA image, and projection of the diffusion measures to the skeleton.

For the statistical comparisons, a permutation-based tool from FSL called *randomise* was used with 5000 permutations following the threshold-free cluster enhancement (TFCE) procedure^41,42^. Besides, to consider statistically significant results within regions, p < 0.05, family-wise error corrected with the TFCE option, and regions of interest (ROI) greater than 30 mm3 were used as threshold. The same basic procedure as in ^27,28^ was followed.

Further, Cohen’s D value was calculated over the ROIs > 2500 mm^3^ to see the variability within the different DTI and AMURA measures. In addition, the full WM Cohen’s D values was obtained to better describe what happened with each measure in the whole brain.

#### 2.4.2. Resampling of diffusion measures

To better understand how the diffusion measures behave in relation to the number of subjects in each group, a resampling experiment was carried out. Thereby, a ROI based analysis was performed to understand how the statistical differences in each region vary depending on the sample size, in comparison with those results obtained in the original TBSS analysis, explained in the previous section.

In this experiment, the number of subjects of each group (EM, CM and HC) was progressively decreased from the original number to 10 subjects in each group, reducing five subjects for each iteration. This process was stopped if no regions with significant differences were found for a particular iteration. For each iteration, 201 different subsamples were randomly obtained following a bootstrapping procedure.

The average value of the measures on the FA-skeleton inside each ROI from the JHU WM atlas was calculated using the 2% and 98% percentiles and compared with an ANOVA test. For the tests with statistically significant differences, post-hoc two-by-two comparisons were carried out, producing a total of 3 comparisons (EM vs. HC, CM vs. HC and CM vs. EM). Finally, the result of the post-hoc test was considered statistically significant in each ROI for p < 0.05 and p < 0.01. No kind of statistical correction was considered for this experiment considering that our purpose was to study the behavior of the different metrics with the sample size.

This methodology was carried out for all the considered measures for both the DTI and AMURA approaches.

#### 2.4.3. Analysis of stability

The coefficient of quartile variation (CQV) was used to measure the stability across groups. The CQV is a measure of homogeneity ^40^ and it was used to assess the inter-subject variability, considering the diverse sample sizes from the analysis described in the previous section. The CQV is one of the most robust statistical measures as it depends on the quartiles, being less sensitive to outliers. Its use is as follows:

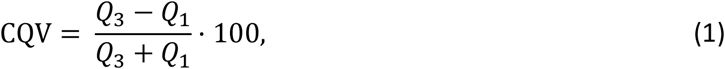

where *Q*_1_ and *Q*_3_ are the first and third quartile, respectively.

The CQV is calculated for each group and ROI, considering as figure of merit the median value of all the CQV of the different 201 subsamples used in this experiment. The 95% confidence interval (95% CI) was set taking the 2.5 and 97.5 percentiles of the whole CQV values for each group of values. This 95% CI was compared between the diverse measures and regions for each sample size.

## 3. RESULTS

The detailed demographic and clinical features of the three groups are shown in Table 1. No statistically significant differences in age or gender were found between the three groups. Patients with CM showed significantly higher duration of migraine, frequency of headache and migraine attacks and medication overuse, and a lower presence of aura.

**Table 1.** Clinical and demographic characteristics of healthy controls (HC), episodic migraine (EM) and chronic migraine (CM).

|  | HC (n=50) | EM (n=51) | CM (n=56) | Statistical test |
| --- | --- | --- | --- | --- |
| Gender, male/female | 11/39<br>(22/78%) | 7/44<br>(14/86%) | 6/50<br>(11/89%) | $\chi^2(2, N = 157) = 2.74$<br>$p = .25^+$ |
| Age (years) | 36.1 ± 13.2 | 36.6 ± 7.9 | 38.1 ± 8.7 | $\chi^2(2) = 2.79, p = .25^+$ |
| Duration of migraine history (years) | | 13.1 ± 10.5 | 19.6 ± 10.4 | $U = 932.5, p = .002^\S$ |
| Time from onset of chronic migraine (months) |  |  | 24.5 ± 32.9 |  |
| Headache frequency (days/month) | | 3.6 ± 1.9 | 23.3 ± 6.3 | $U = 40.0, p < .001^\S$ |
| Migraine frequency (days/month) | | 3.6 ± 1.9 | 13.9 ± 6.9 | $U = 99.5, p < .001^\S$ |
| Medication overuse | | 0 (0%) | 42 (75%) | $p < .001^*$ |
| Aura | | 9 (18%) | 1 (2%) | $p = .006^*$ |

<sup>+</sup>Chi-square test. <sup>†</sup>Kruskal-Wallis test. <sup>§</sup>Mann-Whitney U test. <sup>\*</sup>Fisher's exact test. Data are expressed as means ± SD.

### 3.1. TBSS (original sample)

Using the DTI measures (FA, MD, AD and RD), statistically significant differences between CM and EM patients were observed for two parameters. Patients with CM showed lower AD and MD values than EM in 40 and 38 out of 48 regions from the JHU-WM Atlas, respectively. No statistically significant differences were found using DTI measures between EM and HC or between CM and HC.

For the AMURA metrics, the comparison between patients with EM and HC showed the highest number of parameters with statistically significant differences. Significant lower RTOP, RTAP, qMSD, APA, DiA and ϒ ^1/2^ values in EM compared to HC were found in 41, 39, 43, 27, 29, and 9 ROIs out of 48, respectively. Concerning the comparison between both groups of patients, higher values in CM compared to EM were identified for the RTPP and ϒ ^1/2^ in 4 and 7 regions, respectively.

Figure 2 shows the TBSS results including all the ROIs that presented statistically significant differences together with the FA skeleton, colored in dark blue, for each of the three comparisons. On the one hand, for EM vs. HC and CM vs. HC comparisons, all the AMURA measures which showed significant differences are merged and depicted in the figure, that is, RTOP, RTAP, APA, qMSD, ϒ ^1/2^ and only DiA for EM vs. HC. On the other hand, DTI and AMURA measures can be distinguished in the last CM vs. EM comparison. For DTI, the measures merged depicted are AD and MD, while for AMURA are RTPP and ϒ ^1/2^. As it can be seen, AMURA measures showed differences in group comparisons where the DTI ones did not, as shown in the green circles. A summary with the previous TBSS results regarding the number of ROIs and the group comparisons can be found in Figure 3.

**Figure 2.**
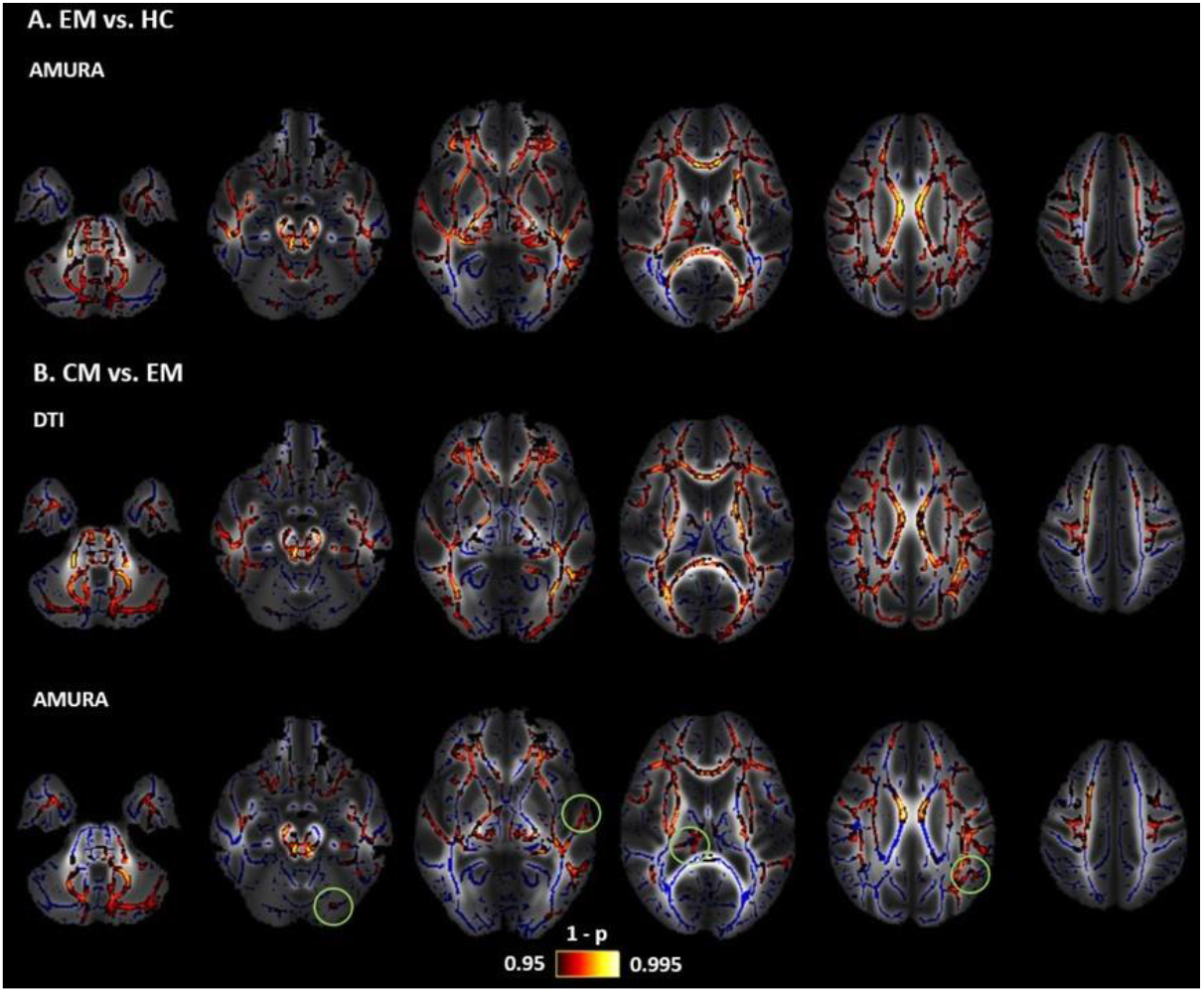
Statistically significant clusters of voxels distinguishing between DTI and AMURA approaches. Mean FA image at the background, FA skeleton coloured in blue and significant ROIs coloured in red-yellow. **A)** Episodic Migraine (EM) vs. Healthy Controls (HC): merged AMURA measures (RTOP, RTAP, APA, qMSD, ϒ ^1/2^ and DiA). **B)** CM vs. EM: merged DTI (AD and MD) and AMURA (RTPP, and ϒ ^1/2^) measures. DTI measures do not detect any significant ROI either in EM vs. HC nor CM vs. HC. Green circles showed the areas where AMURA measures showed differences in group comparisons where the DTI ones did not.

**Figure 3.**
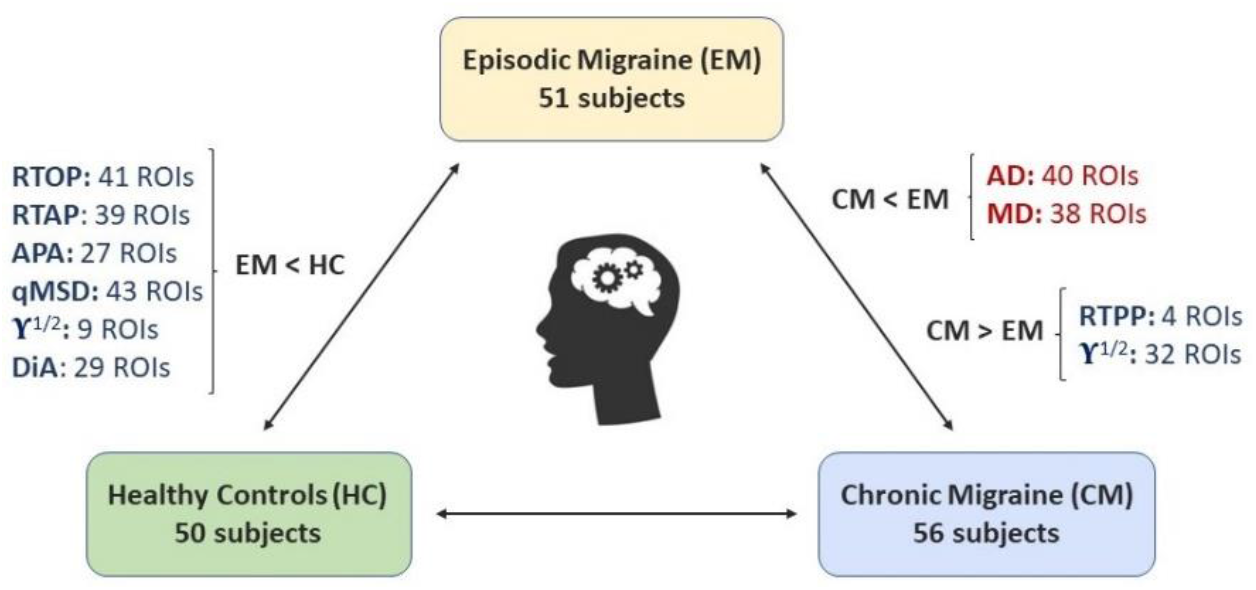
Summary of the main tract-based spatial statistics (TBSS) results with DTI (in red) and AMURA measures (in dark blue).

Detailed results are available in Appendix C.

The effect size for these comparisons varied among the different diffusion measures. Figure 4 shows the absolute value of Cohen’s D in ROIs greater than 2500 mm^3^, and for the three group comparisons.

**Figure 4.**
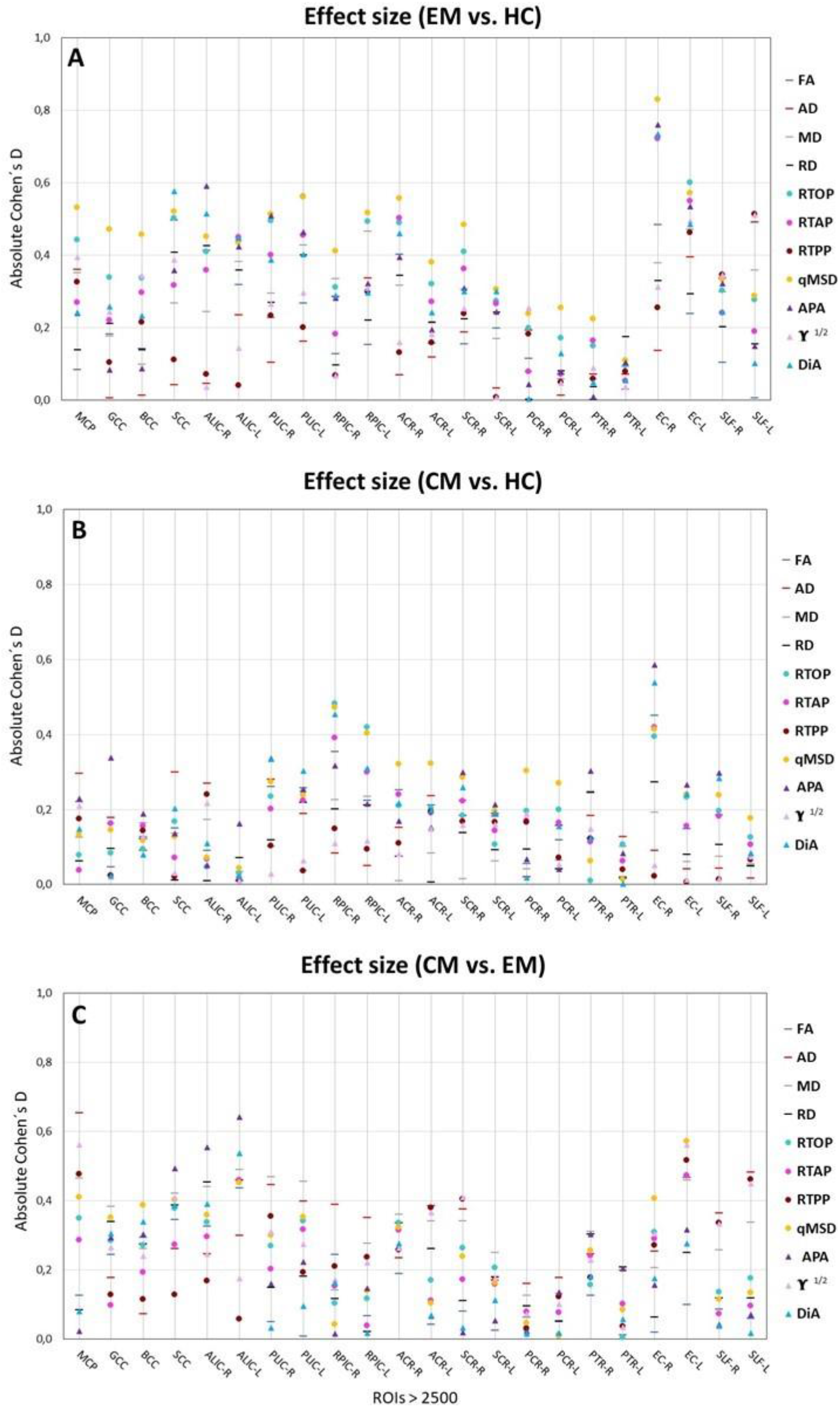
Absolute Cohen’s D represents the effect of the DTI (small line) and AMURA (circles) measures, which have shown significant differences. ROIs greater than 2500 mm^3^ according to the JHU WM Atlas were considered. **A)** Episodic Migraine (EM) vs. Healthy Controls (HC). **B)** Chronic Migraine (CM) vs. HC. **C**) CM vs. EM.

It can be observed (Figure 4A) that for the comparison between EM and HC, the qMSD showed the largest effect sizes as measured by larger Cohen’s D values. For instance, regions such as the right external capsule (EC-R), the right anterior corona radiata (ACR-R) or the right posterior limb of internal capsule (PLIC-R) for the qMSD measure showed an absolute value of Cohen’s D higher than 0.55, indicating a moderate-large effect size (the threshold for medium effect size is 0.5 and for large effect size is 0.8). In addition, these regions presented significant differences with the TBSS approach.

Regarding the comparison between CM and HC (Figure 4B), the right external capsule (EC-R) for the APA and the DiA reached absolute values of Cohen’s D higher than 0.4, showing at the same time significant differences with TBSS.

Finally, in the comparison between CM and EM (Figure 4C), the middle cerebral peduncle (MCP) for the AD and the left anterior limb of the internal capsule (ALIC-L) for the APA showed Cohen’s D value higher than 0.6, indicating a moderate-large effect size while providing statistically significant differences with TBSS.

It is also interesting to analyze the behavior of each measure over the whole WM. Figure 5 shows the absolute Cohen’s D in the whole WM for each measure, together with the total number of ROIs with significant differences obtained with the TBSS approach. The biggest effect sizes were obtained for the comparison between EM vs. HC. Coherently, this comparison also produced the highest number of ROIs with significant differences when using TBSS. For instance, the qMSD or the RTOP measures reached absolute Cohen’s D values greater than 0.6, and more than 40 ROIs with significant differences in TBSS. On the other hand, the comparison between CM and HC presented the lowest Cohen’s D values. Regarding the comparison between CM and EM, the AD, MD and ϒ ^1/2^ measures have more than 30 significant ROIs and Cohen’s D values greater than 0.45.

**Figure 5.**
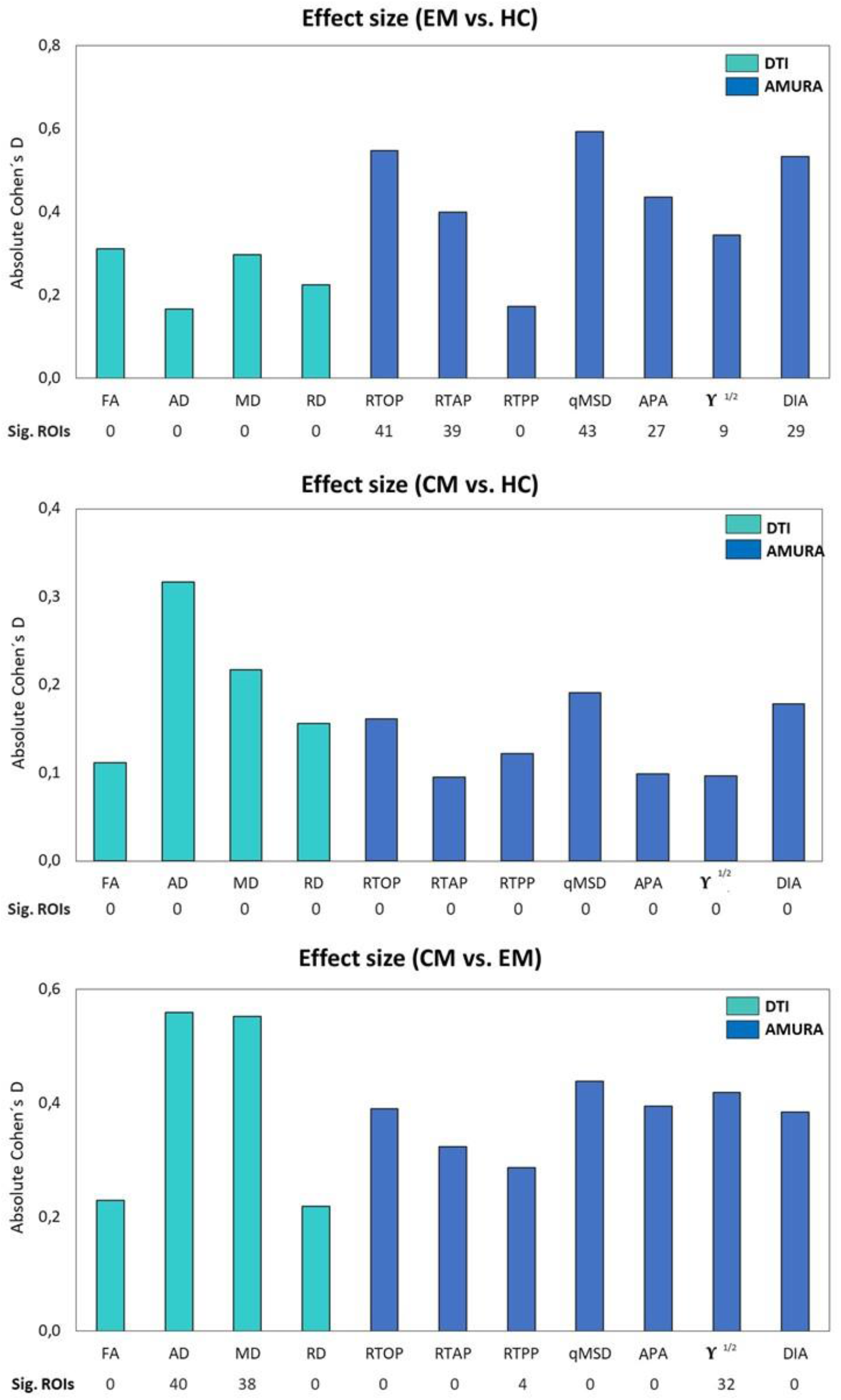
Absolute value of effect sizes for the three group comparisons in the Full WM: Episodic Migraine (EM) vs. Healthy Controls (HC), Chronic Migraine (CM) vs. HC and CM vs. EM. DTI and AMURA measures are depicted. For each measure, the total number of ROIs that presented statistically significant differences obtained with the TBSS approach is also noted, to relate both studies.

### 3.2. Resampling of diffusion measures

Figure 6 shows the effects of changing the sample size for the different DTI and AMURA-based measures. Among the DTI measures, results with MD showed a relatively high number of ROIs with statistically significant differences using bigger samples sizes, as can be seen in Figure 6C. However, the number of significant ROIs drastically decreased for a group sample of 40. In addition, few ROIs with statistically significant differences were found for the rest of DTI measures and for the other two group comparisons, in any sample size, which made the assessment of the relationship between DTI measures and sample size unfeasible.

**Figure 6.**
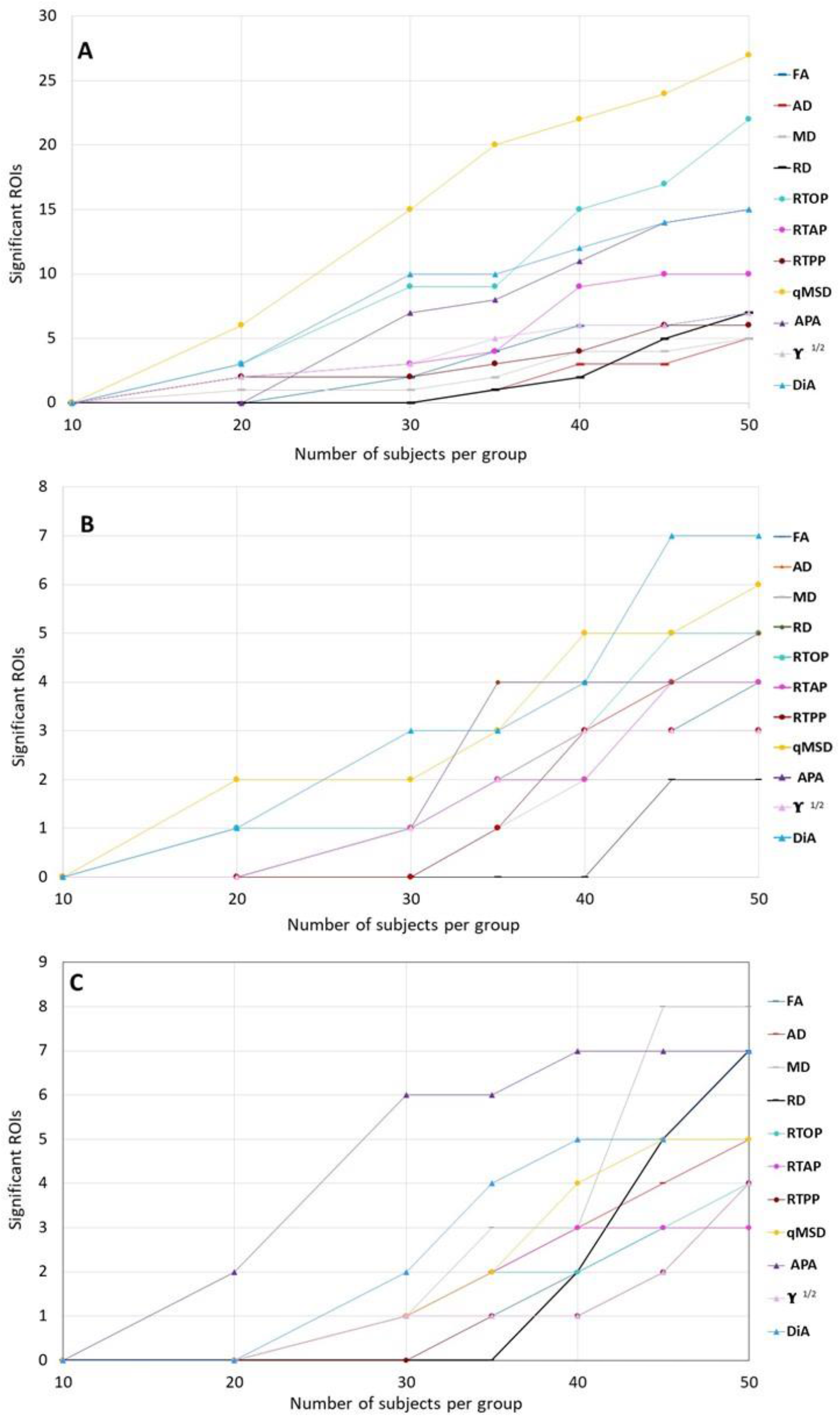
Significant ROIs by the resampling of diffusion measures reducing the number of subjects per group. **A)** Episodic Migraine (EM) vs. Healthy Controls (HC). **B)** Chronic Migraine (CM) vs. HC. **C)** CM vs. EM. DTI (triangles) and AMURA (circles) measures are shown in the graph. No statistical correction was considered.

Considering the results with *p* < 0.05, Figure 6 shows a stable behavior of AMURA measures in relation to the sample size, which can be understood as a linear dependence between the group sample size and the number of statistically significant ROIs. In Figure 6A, this behavior can be better understood and interpreted in measures such as qMSD, which is the most robust one in the comparison between EM and HC. Furthermore, RTOP is also the most robust measure in the CM vs HC comparison. Notice that most of AMURA measures reached the lack of statistical significance ROIs for a group sample of 10.

Regarding the results with *p* < 0.01 (Appendix D), similar trends were observed compared to the assessment with *p* < 0.05, being the qMSD or RTOP the parameters with higher number of ROIs with statistically significant differences when reducing the sample size.

Figure 7 shows a p-value scheme for the 48 JHU WM atlas ROIs considered for the 201 different subsamples of the total number of subjects per group, where two-by-two group comparisons using DTI and AMURA are depicted. Most of the significant ROIs obtained using the traditional DTI measures were also present with the AMURA ones. However, as it can be seen in the figure, AMURA measures revealed additional ROIs with statistically significant differences in comparison with DTI.

**Figure 7.**
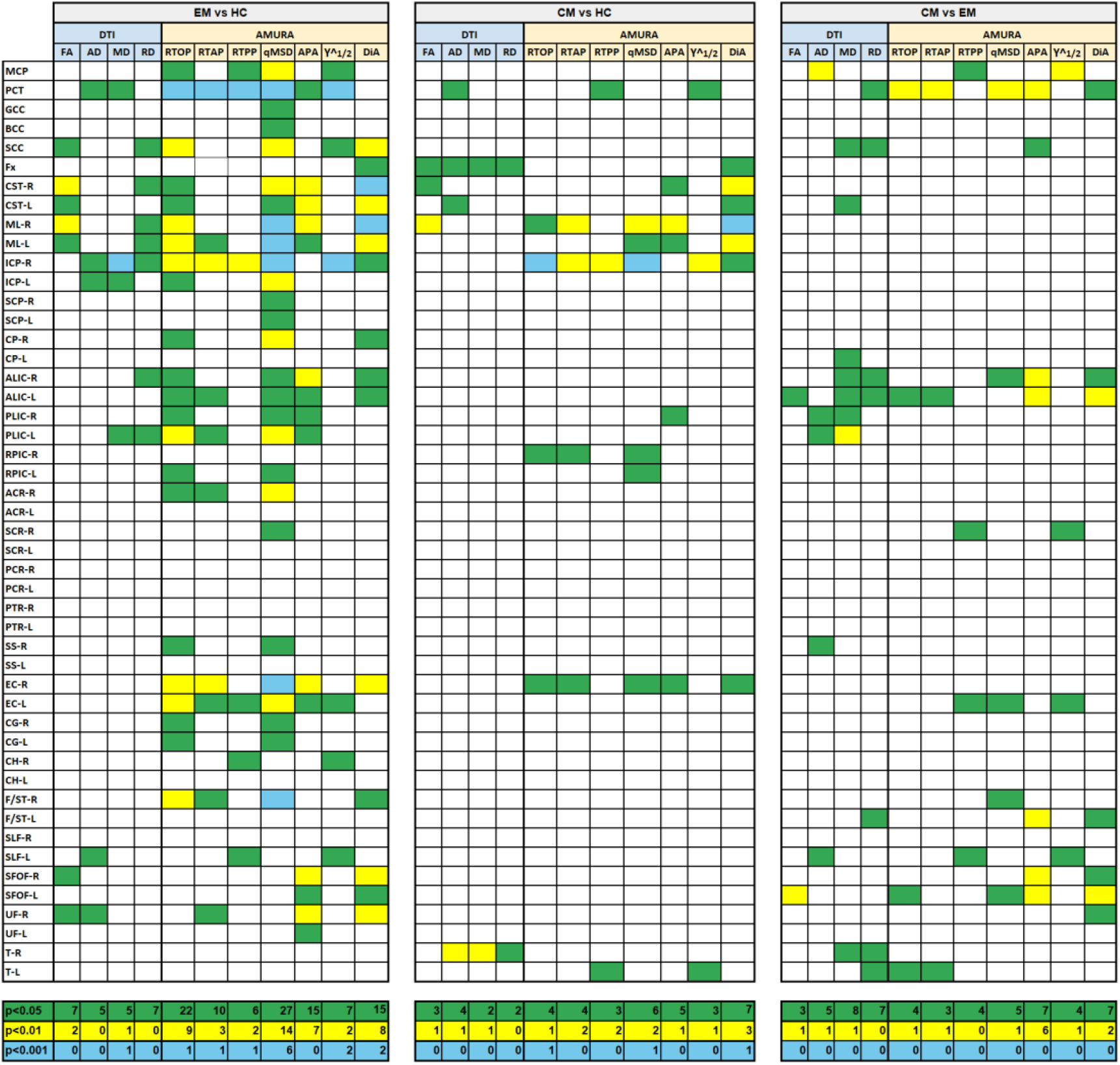
Episodic Migraine (EM) vs. Healthy controls (HC), Chronic Migraine (CM) vs. HC and CM vs. EM. Two-sample t-tests for DTI and AMURA measures and each of the ROIs defined by the JHU WM atlas. The p-values represent the probability that a certain measure has identical means for both groups. ROIs exhibiting differences with statistical significance above 95% (*p <* 0.05) are highlighted in green, above 99% (*p <* 0.01) in yellow and above 99.9% (*p* < 0.001) in blue. In the bottom of the figure, the number of regions for each measure and p-value are also shown. Notice that blue regions are also yellow and green ones, meaning the significant ROIs for *p <* 0.01 and *p <* 0.05.

### 3.3. Analysis of stability

Figure 8 depicts the average values of CQV for all the DTI and AMURA-based diffusion measures. The measures with the highest stability (lowest CQV) were the RTPP and the APA, with an approximate average CQV of 2% considering all the regions. Other measures with relatively high stability were the three DTI measures (FA, MD and AD), ϒ ^1/2^ and DIA, with CQV average values between 2% and 5%. The remaining DTI and AMURA descriptors (RD, RTAP and RTOP), presented a moderate-high stability, with CQV average values between 5% and 10%. The descriptor with the lowest stability was the qMSD, with CQV average values between 15% and 20%.

**Figure 8.**
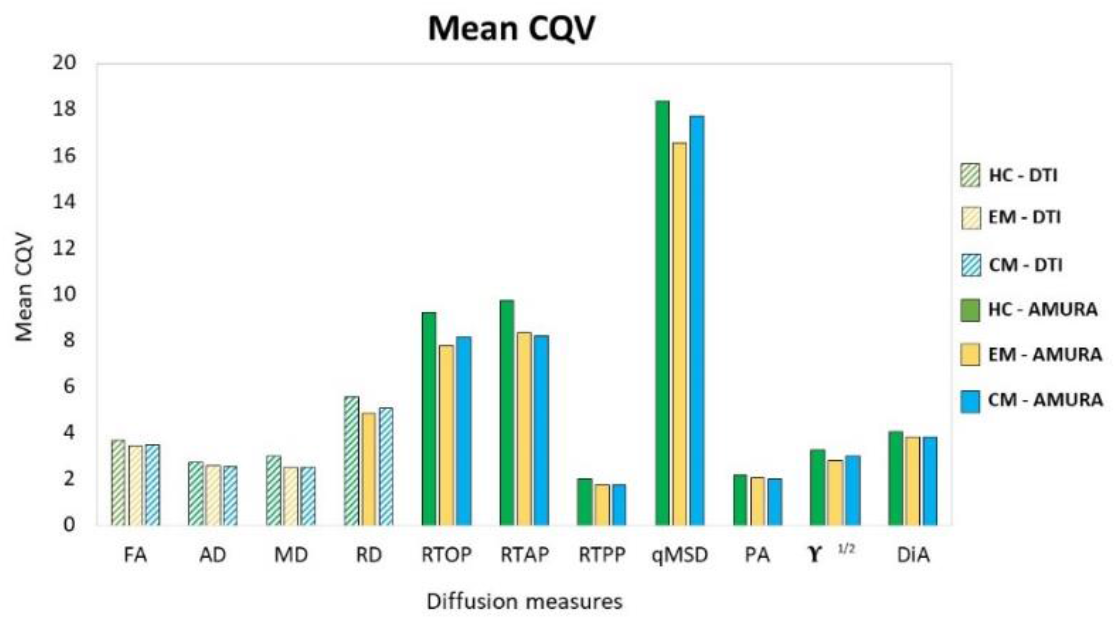
Mean CQV for each group of study considering the 48 ROIs of JHU-WM atlas. Healthy Controls (HC), Episodic Migraine (EM) and Chronic Migraine (CM). DTI and AMURA measures are shown. The measures with the higher stability have lower CQV.

Regarding the comparisons between the three groups of interest, after reducing the group sample size to 45 subjects, the assessment of the CQV 95% CI showed that the HC presented a general higher variability than patients with EM and CM. The parameters with a higher number of regions with statistically significant differences between HC and migraine patients according to the 95% CI were the three AMURA measures (RTOP, RTPP and RTAP) and the MD, with 14-22 regions presenting differences. Additionally, in the comparison between HC and CM, the CQV of APA or ϒ^1/2^ were significantly higher in HC than CM in 13 regions. The number of regions with CQV differences between CM and EM was lower compared to the comparison between HC and the patient groups. FA and MD were the descriptors with a higher number of regions (nine) that showed higher variance in EM compared to CM, and MD was also the parameter with more regions (eight) with significantly higher variance in CM.

## 4. DISCUSSION AND CONCLUSIONS

Advanced diffusion descriptors such as RTOP, RTAP and APA have shown to be useful for the analysis of the WM of the brain ^24,27,28^. However, their conventional calculation requires acquisition protocols including several b-values, a high number of diffusion gradient directions and very long processing times. This makes them unfeasible for their use in clinical practice or in many commercial MRI scanners. Besides, the use of these metrics in retrospective studies is usually impossible since the acquisition protocols do not allow for it.

AMURA was proposed to allow the estimation of *apparent* versions of these advanced diffusion measures from reduced acquisitions ^24–26^. It provides a fast and straightforward method to compute them from a single shell and very short processing times. Metrics calculated with AMURA have shown a high correlation with measures calculated using a multishell approach, such as MAP-MRI^13^, MAPL^50^ or MiSFIT^14^, for high b-values (at least 2000 s/mm^2^). For lower values, these measures show a weaker correlation since the underlying features measured are better visible at higher b-values ^24,25^. However, we hypothesized that AMURA metrics can still provide useful information at lower b-values, complementary to that obtained from DTI-based measures. This paper focuses precisely on that hypothesis and tries to elucidate whether AMURA-based measures obtained from standard DTI-type acquisitions are useful in group studies.

To that end, we have resorted to migraine as our target pathology, because of several reasons. First, diffusion MRI studies in the literature show that differences between patients and HC, or between different groups of patients (EM vs. CM) are subtle, as studies using small sample sizes have often reported no differences and even contradictory findings have been published ^30,41–46^.

To study the viability of AMURA-based measures, a conventional TBSS analysis together with the assessment of the behavior of the diverse measures from reduces sample sizes and of the stability were carried out.

Indeed, results showed that AMURA measures obtained from DTI-type acquisitions were able to successfully find statistically significant differences between the three groups under study (HC, EM and CM), including differences that were not detected using DTI-based measures. Although AMURA showed additional differences between groups in a preliminary previous study ^27^, the magnitude of the additional differences, particularly those between EM and HC, was unexpected.

As depicted in Figure 3, a summary of the significant differences between groups detected by AMURA-based and DTI-based measures reveals their complementary nature. Indeed, DTI-based AD and MD showed a good performance for the comparison between EM and CM, while some of the AMURA-based measures were also able to find statistical differences. Finally, AMURA-based measures showed their sensitivity for the comparison between EM and HC, detecting differences with most measures and in a high number of regions. Many of those regions include the pain matrix and thalamic-cortical connections, involved in pathophysiology of migraine and other chronic pain syndromes.

Although obtained with a different experiment, results in Figure 7 also highlight the complementary nature of AMURA-based measures with respect to DTI-based ones. While AMURA-based measures detected differences in several regions for which DTI-based measures also did (such as MCP, SCC and ALIC), they also found differences in regions where DTI-based measures did not (PCT, EC-L, F/ST-L and T-L, among others). The opposite is also true for some other WM regions (CST-L, CP-L, SS-R and T-R).

The sensitivity of AMURA-based measures was further analyzed by studying the effect size found in the different comparisons between groups. A classical method to determine the magnitude of the differences between groups is Cohen’s D, which considers the variability of the sample in relation to the average value. As illustrated in Figures 4 and 5, DTI-based and AMURA-based measures showed comparable effect sizes for the CM-EM and CM-HC comparisons. In the first case, DTI-based AD and MD reached medium effect sizes (0.5) (for the whole WM), while Cohen’s D for FA or RD barely exceeded small effect size threshold (0.2). For this comparison, Cohen’s D for AMURA-based measures varied between the small and the medium effect thresholds. For the comparison between CM and EM, however, Cohen’s D values were notably lower for all measures, barely reaching 0.3 for DTI-based AD. Finally, regarding the comparison between EM and HC, while DTI-based FA and MD reached Cohen’s D values around 0.3, AMURA-based RTOP, qMSD and DIA reached values over 0.5. These differences in effect sizes among different measures and different group comparisons offer a good explanation for the results shown in Figures 2 and 7 and summarized in Figure 3.

Whereas it may be tempting to think about EM and CM as different degrees of the same pathological process, recent results ^30,41^ support the hypothesis of EM and CM being different entities at the microstructural level, each accompanied by different changes in the WM. Following this hypothesis, DTI-based measures seem well-fitted to detect WM changes in CM, while AMURA-based methods perform remarkably well for the changes that occur in EM. Although the interpretation of changes in DTI or AMURA-based measures is not straightforward, results suggest that WM changes in EM with respect to HC (specifically, lower RTOP and RTAP) might be related to changes in the transverse diffusivity, while changes in CM with respect to EM (such as higher RTPP and lower AD) might be more related to changes in the diffusivity in the axonal or main direction.

Considering the difficulty to obtain large sample sizes in group studies, it is important to assess the behavior of the diverse diffusion measures when the number of subjects per group is reduced. As depicted in Figure 6, both DTI-based and AMURA-based measures share the expected trend, meaning that the number of ROIs with statistically significant differences decreases as the sample size is reduced. Remarkably, DTI-based MD shows a more pronounced reduction in the number of significant ROIs is reduced for the comparison between CM and EM.

The assessment of the stability provides another interesting perspective for the evaluation and comparison between different diffusion measures. The diffusion measures that showed higher stability (lower CQV) were AMURA-based APA and RTPP and the DTI-based measures, while AMURA-based qMSD seems to present low stability. This high variability was expected, since qMSD is a quadratic measure, so it must show a greater range of variability. Interestingly, it presented a relatively high number of regions with statistically significant differences in the comparisons of both migraine groups against controls for diverse sample sizes despite their low stability. Therefore, the results of this study suggest that qMSD is able to characterize specific microstructural properties that are particularly difficult to find with other parameters.

Moreover, as it has been suggested previously in this section, differences between both groups of patients with migraine and controls may be qualitatively distinct compared to the differences between CM and EM. Furthermore, qMSD is especially sensitive to short diffusion time scales ^16^.

It is important to note that the AMURA-based measures employed in this paper must be considered as *apparent* values at a given b-value, and their interpretation in terms of the microstructure properties may be different from that of the original EAP-based diffusion measures. Although the relationship between AMURA-based measures and their original counterparts deserves further study, in this paper we deliberately chose not to pursue this comparison to focus on the viability of AMURA-based measures to complement DTI in scenarios where EAP-based measures cannot be obtained.

This study presents limitations that must be pointed out. First, the pathophysiological interpretation of the different trends of the AMURA-based measures is not totally clear, so a description of the microstructural properties according to the values of each measure cannot be provided. As mentioned previously, the apparent nature of AMURA-based measures and their complex relationship with the original EAP-based measures prevent the direct adoption of interpretations from those EAP-based measures. Microstructural studies like those conducted for DTI-based measures ^48,49^ are needed to fully understand the results obtained with AMURA.

Furthermore, the results obtained in this study cannot be directly translated to other pathologies affecting the WM of the brain. Even though AMURA can be expected to be a useful information to detect differences in group studies targeting other diseases, further research is needed to confirm that.

In conclusion, this study showed that the new AMURA-based measures can be easily integrated in group studies using single-shell dMRI acquisition protocols, and they can reveal WM changes that may remain hidden with traditional DT-based measures. The wide variety of AMURA, a fast and relatively simple approach, provides measures that allow to extract values that are able to find differences between groups for restricted sample sizes and dMRI acquisition protocols.

## ACKNOWLEDGMENTS

This work was supported by Ministerio de Ciencia e Innovación of Spain with research grant RTI 2018-094569-B-I00 and Gerencia Regional de Salud de Castilla y León with research grant GRS 1727/A/18. Á.P.-G. was supported by the European Social Fund (NextGenerationEU).

## Abbreviations

AD: axial diffusivity
ALIC-L: left anterior limb of the internal capsule
AMURA: Apparent Measures Using Reduced Acquisitions
APA: apparent propagator anisotropy
CI: confidence interval
CM: chronic migraine
CQV: coefficient of quartile variation
DiA: Diffusion Anisotropy
DDT: divergence from Diffusion Tensor
DTI: diffusion tensor imaging
DWI: diffusion-weighted image
EAP: Ensemble Average Propagator
EC: external capsule
EC-R: right external capulse
EM: episodic migraine
FA: fractional anisotropy
HC: healthy controls
MCP: middle cerebral peduncle
MD: mean diffusivity
qMSD: Q-Space Mean Square Displacement
RPIC-L: right superior corona radiata
RTAP: return-to-axis probabilities
RTOP: return-to-origin
RTPP: return-to-plane
SCR-R: right superior corona radiata
SH: Spherical Harmonics
TE: echo time
TR: repetition time
WM: white matter

## Appendix A. Graphical abstract figure

Figure A1 shows a graphical abstract of the calculation of AMURA methodology. It is briefly summarized.

**Figure A1.**
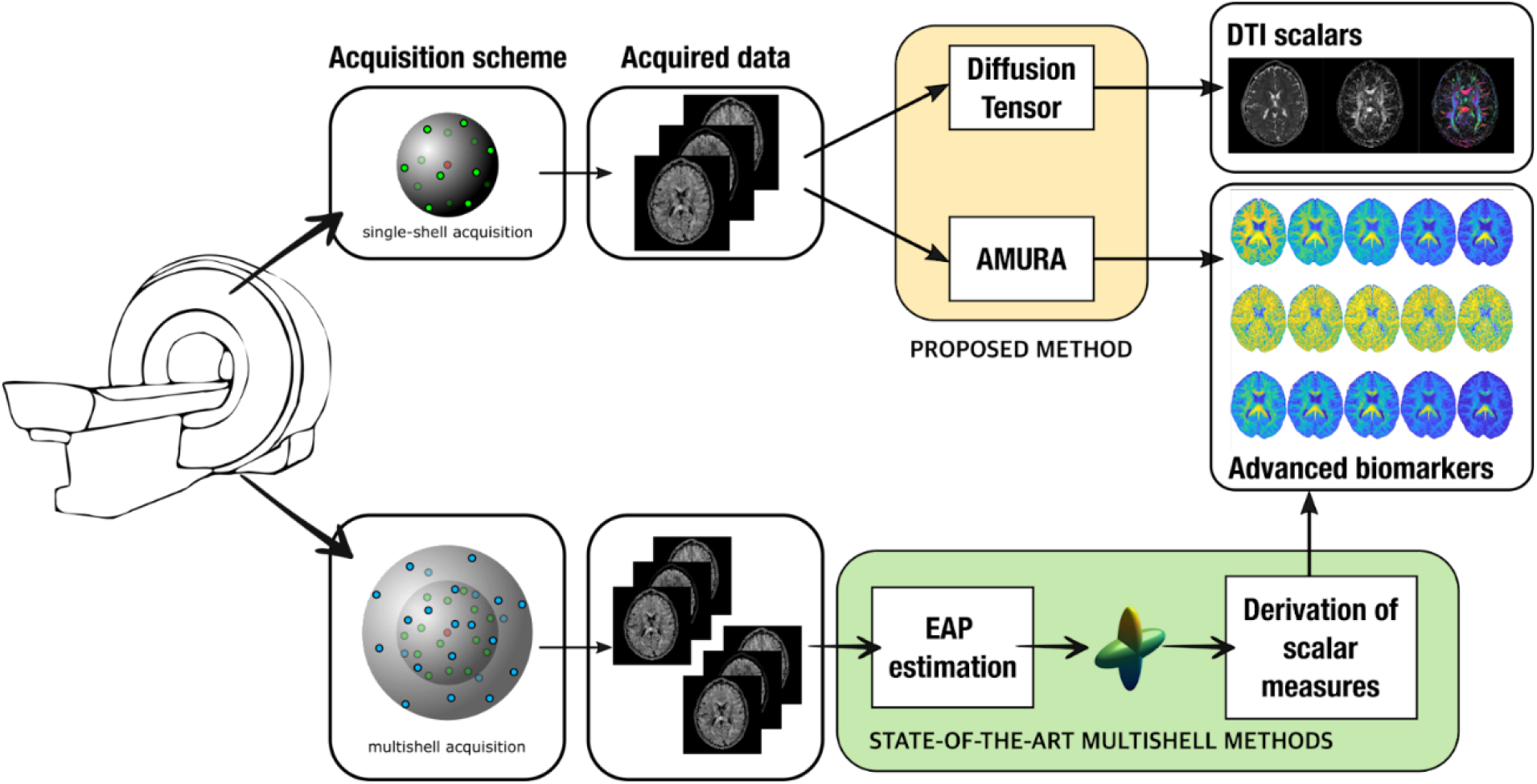
Graphical abstract of the calculation of AMURA.

### B. Definition of AMURA metrics used in the study

AMURA allows the estimation of the EAP-related scalars without the explicit calculation of the EAP, using a lower number of samples from a single-shell acquisition scheme^24–26^. AMURA considers that, if the amount of data is reduced, a restricted diffusion model consistent with single-shell acquisitions must be assumed: the ADC *D*(***q***) does not depend on the radial direction (i.e., on the magnitude of the q-vector) within the range of b-values probed, so that *D*(***q***) = *D*(***u***) where ***u*** is a unit direction in space where ‖***u***‖ = 1 and ***q*** = *q****u***. This way, assuming a general Gaussian diffusion profile, the normalized magnitude image provided by the MRI scanner, *E*(***q***), becomes:

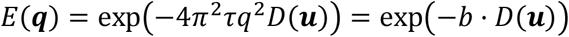

From this equation, AMURA proposed a particular implementation of scalar measures. Since the mono-exponential model only holds within a limited range around the measured b-value, the measures derived this way must be seen as *apparent* values at a given b-value, related to the original ones but dependent on the selected shell. The AMURA metrics used in the paper are the following ones:

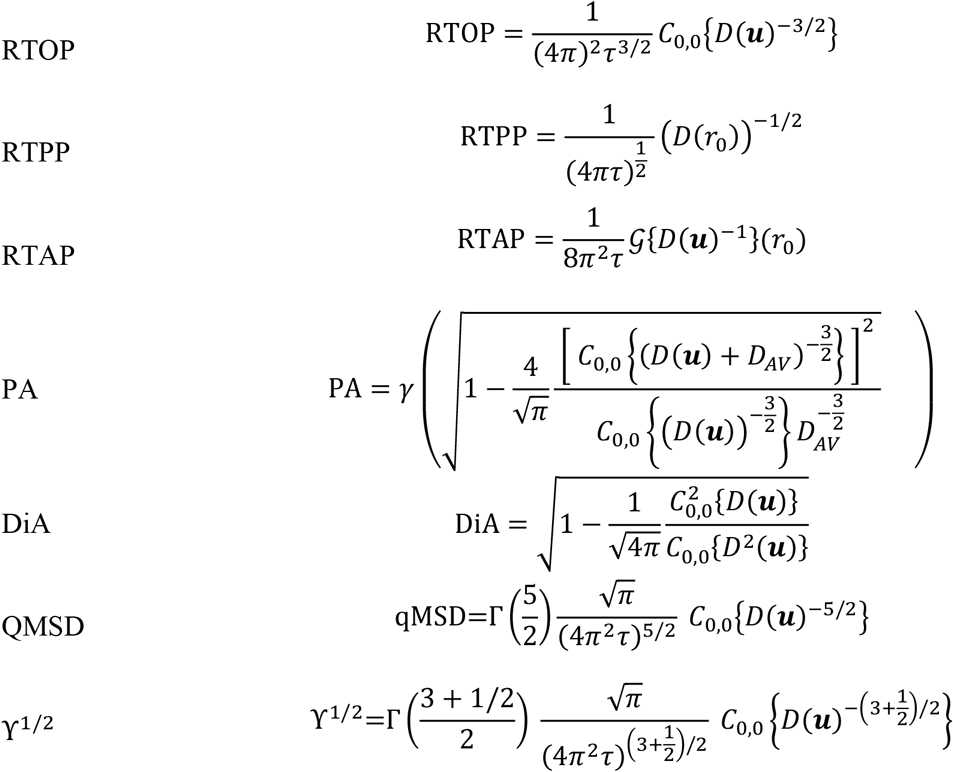

Notation:

- 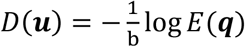
- *C*_0,0_{*H*(***u***)} DC component of signal *H*(***u***) calculated using the zero-th order coefficient of a Spherical Harmonics (SH) expansion.
- *r*_0_ direction of maximal diffusion.
- 𝒢{*H*(***u***)}(*r*_0_) Funk-Radon Transform of *H*(***u***) evaluated in the direction of maximal diffusion *r*_0_
- 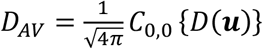 average diffusivity.

## Appendix C. TBSS results

### C.1. DTI measures

TBSS results for AD and MD are shown in Figure C1 and Figure C2, respectively. In addition, the statistically significant ROIs, according to the JHU-WM atlas, for each measure are included in Table C1 and Table C2 together with the minimum p-value obtained and the volume of each region.

**Figure C1.**
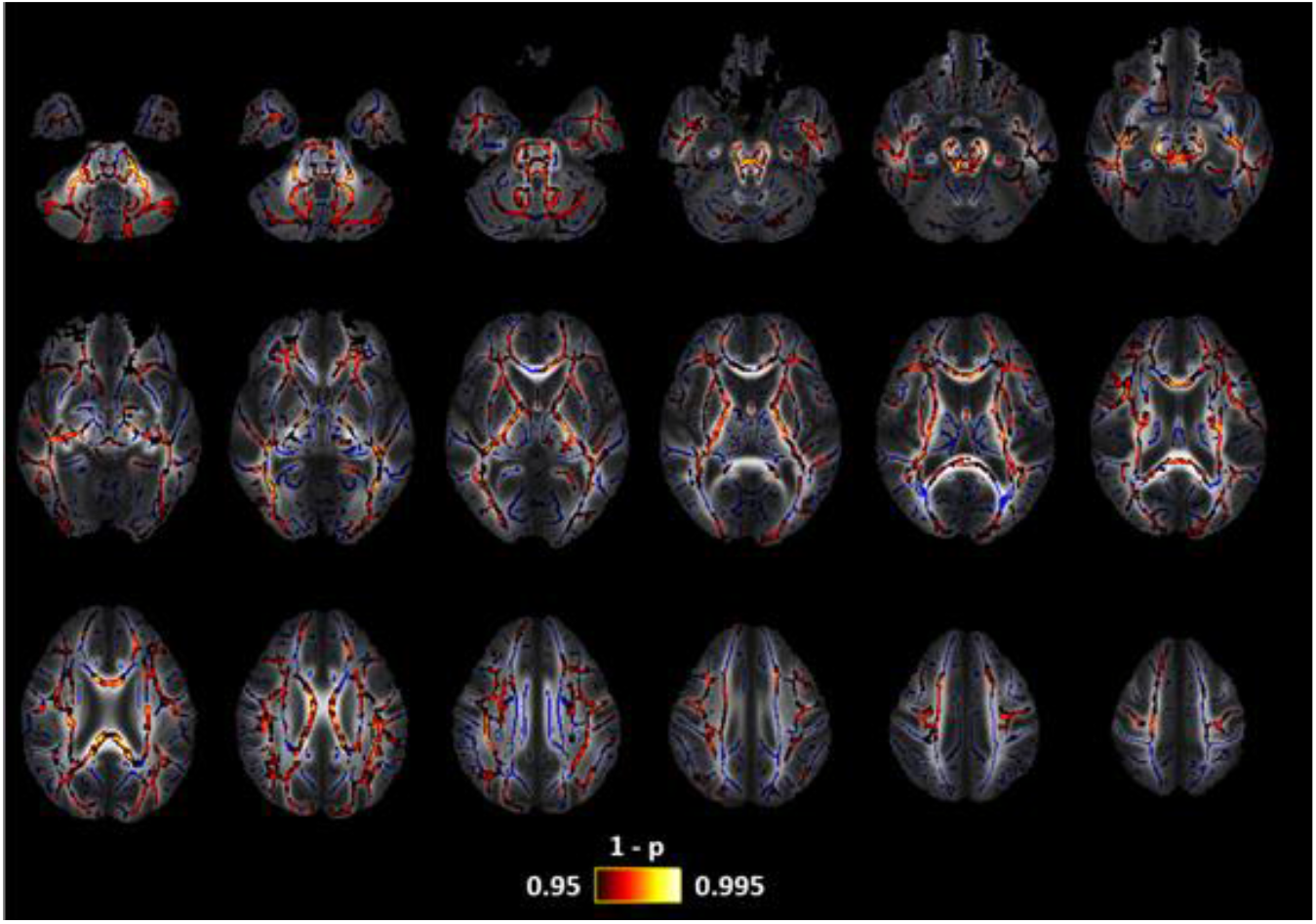
Axial Diffusivity (AD) changes in patients with EM in comparison with CM. Lower AD values were found in CM. The white matter skeleton is shown in blue and voxels with significant differences in red-yellow. The color bar shows the 1-p values (family-wise error corrected).

**Figure C2.**
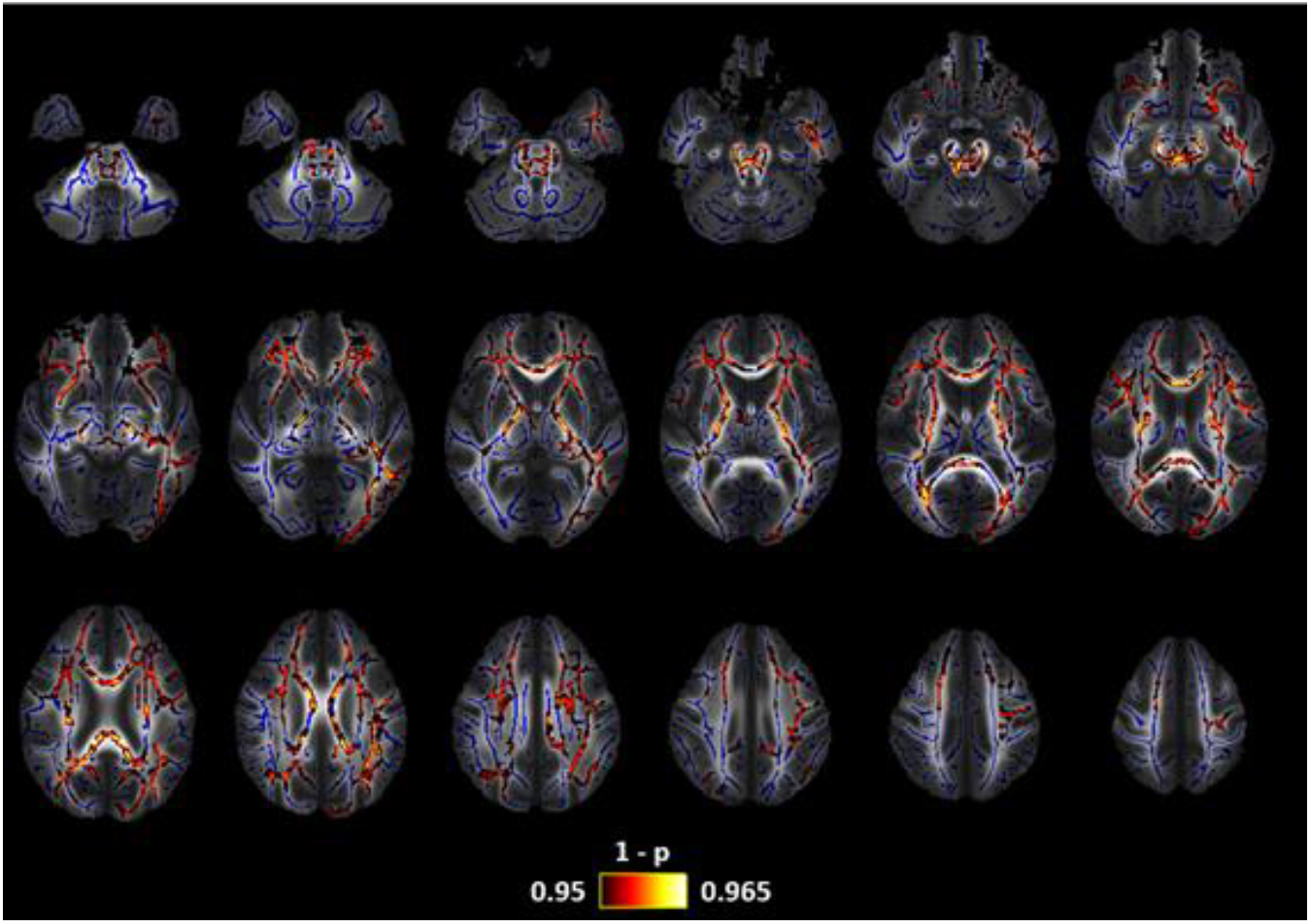
Mean Diffusivity (MD) changes in patients with EM in comparison with CM. Lower MD values were found in CM. The white matter skeleton is shown in blue and voxels with significant differences in red-yellow. The color bar shows the 1-p values (family-wise error corrected).

**Table C1.** White matter regions from the ICBM-DTI-81 White Matter Atlas for which significant decreased AD values were found in CM compared to EM.

| White Matter Region | Minimum *p*-Value (FWE-Corrected) | Volume (mm<sup>3</sup>) |
| --- | --- | --- |
| Middle cerebellar Peduncle | 0.004 | 2175 |
| Superior cerebellar peduncle R/L | 0.0095/0.0095 | 142/122 |
| Inferior cerebellar peduncle R/L | 0.0094/0.005 | 67/74 |
| Superior longitudinal fasciculus R/L | 0.011/0.010 | 871/859 |
| Genu of corpus callosum | 0.012 | 435 |
| Body of corpus callosum | 0.013 | 856 |
| Splenium of corpus callosum | 0.016 | 785 |
| Anterior corona radiata R/L | 0.014/0.010 | 558/707 |
| Superior corona radiata R/L | 0.011/0.009 | 786/446 |
| Posterior corona radiata R/L | 0.013/0.009 | 148/226 |
| External capsule R/L | 0.010/0.009 | 437/541 |
| Posterior limb of internal capsule R/L | 0.011/0.009 | 598/577 |
| Retrolenticular part of internal capsule R/L | 0.012/0.009 | 443/334 |
| Anterior limb of internal capsule R/L | 0.011/0.010 | 270/268 |
| Sagittal Stratum R/L | 0.012/0.009 | 460/356 |
| Posterior thalamic radiation R/L | 0.013/0.010 | 342/253 |
| Cerebral peduncle R/L | 0.008/0.009 | 227/263 |
| Corticospinal tract R/L | 0.008/0.009 | 110/169 |
| Medial lemniscus R/L | 0.009/0.008 | 89/114 |
| Pontine crossing tract | 0.008 | 77 |
| Fornix (cres) R/L | 0.013/0.010 | 91/57 |
| Fornix (body and column) | 0.036 | 44 |
| Cingulum (hippocampus) L | 0.0176 | 169 |
| Tapetum R | 0.013 | 34 |

**Table C2.** White matter regions from the ICBM-DTI-81 White Matter Atlas for which significant decreased MD values were found in CM compared to EM.

| White Matter Region | Minimum *p*-Value (FWE-Corrected) | Volume (mm<sup>3</sup>) |
| --- | --- | --- |
| Middle cerebellar Peduncle | 0.041 | 113 |
| Superior cerebellar peduncle R/L | 0.042/0.043 | 73/32 |
| Superior fronto-occipital fasciculus R/L | 0.038/0.037 | 49/64 |
| Superior longitudinal fasciculus R/L | 0.038/0.035 | 99/714 |
| Genu of corpus callosum | 0.041 | 782 |
| Body of corpus callosum | 0.040 | 948 |
| Splenium of corpus callosum | 0.038 | 966 |
| Anterior corona radiata R/L | 0.038/0.036 | 782/642 |
| Superior corona radiata R/L | 0.038/0.037 | 661/453 |
| Posterior corona radiata R/L | 0.038/0.035 | 100/58 |
| External capsule R/L | 0.040/0.036 | 340/444 |
| Posterior limb of internal capsule R/L | 0.037/0.038 | 509/507 |
| Retrolenticular part of internal capsule R/L | 0.046/0.041 | 113/202 |
| Anterior limb of internal capsule R/L | 0.038/0.036 | 275/363 |
| Sagittal Stratum L | 0.040 | 229 |
| Posterior thalamic radiation R/L | 0.038/0.035 | 136/198 |
| Cerebral peduncle R/L | 0.044/0.040 | 165/208 |
| Corticospinal tract R/L | 0.044/0.040 | 95/144 |
| Medial lemniscus R/L | 0.043/0.044 | 56/47 |
| Pontine crossing tract | 0.043 | 130 |
| Fornix (cres) R/L | 0.046 | 122 |
| Cingulum (cingulate gyrus) L | 0.040 | 74 |
| Tapetum R | 0.040 | 35 |

### C.2. AMURA measures

TBSS results for AMURA measures are displayed as follows: RTOP, Figure C3 and Table C3; RTAP, Figure C4 and Table C4; RTPP, Figure C5 and Table C5; qMSD, Figure C6 and Table C6; APA, Figure C7 and Table C7; **ϒ**^1/2^, Figure C8 and Table C8; DiA, Figure C9 and Table C9.

**Figure C3.**
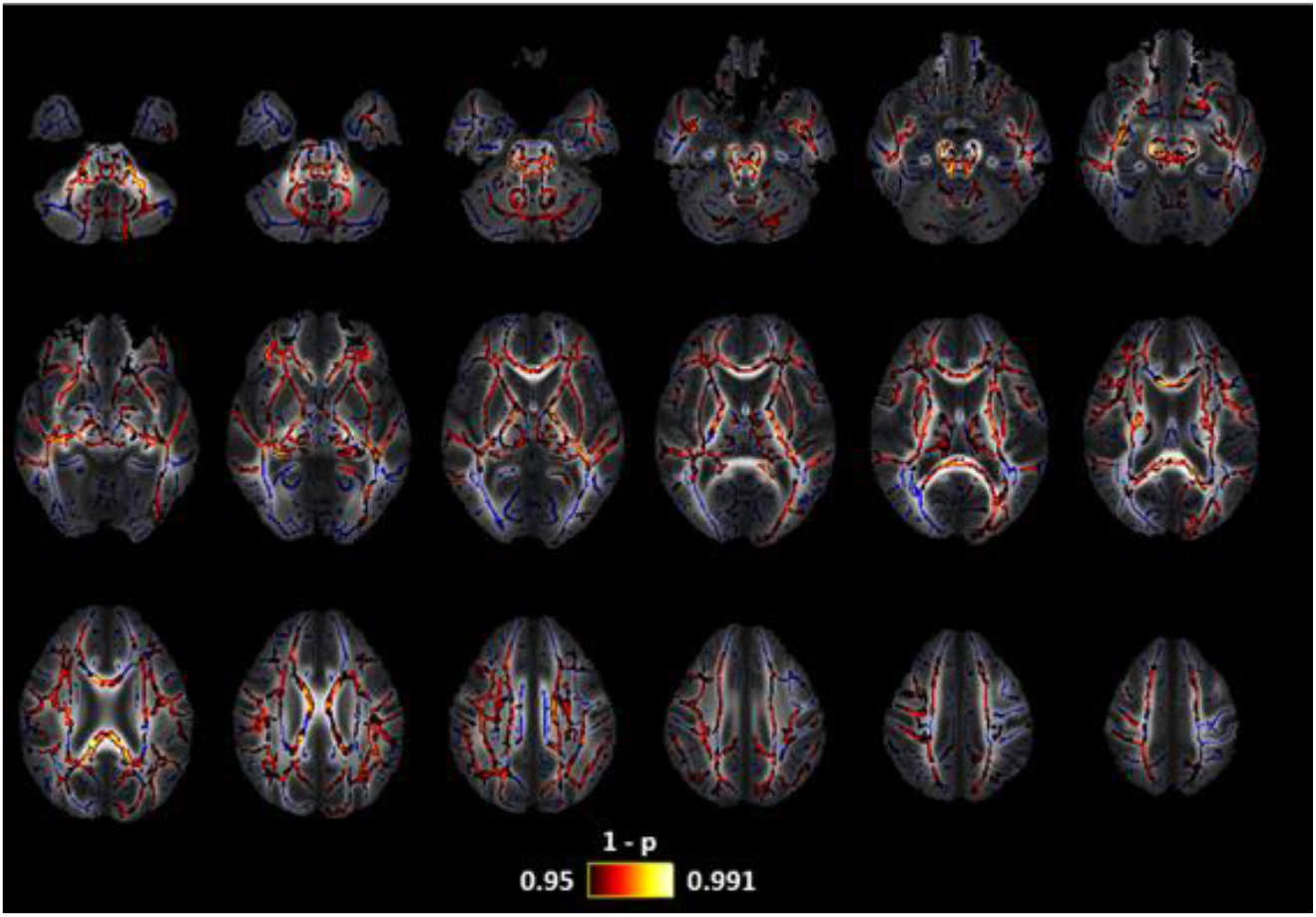
Return-to-origin (RTOP) changes in patients with EM in comparison with HC. Lower RTOP values were found in EM. The white matter skeleton is shown in blue and voxels with significant differences in red-yellow. The color bar shows the 1-p values (family-wise error corrected).

**Table C3.** White matter regions from the ICBM-DTI-81 White Matter Atlas for which significantly decreased RTOP values were found in HC compared to EM.

| White Matter Region | Minimum *p*-Value (FWE-Corrected) | Volume (mm<sup>3</sup>) |
| --- | --- | --- |
| Middle cerebellar Peduncle | 0.0086 | 1663 |
| Superior cerebellar peduncle R/L | 0.086/0.0088 | 134/107 |
| Inferior cerebellar peduncle R/L | 0.0091/0.0099 | 132/93 |
| Superior fronto-occipital fasciculus R/L | 0.009/0.010 | 57/42 |
| Superior longitudinal fasciculus R/L | 0.012/0.010 | 858/705 |
| Genu of corpus callosum | 0.0094 | 801 |
| Body of corpus callosum | 0.012 | 1436 |
| Splenium of corpus callosum | 0.019 | 1592 |
| Anterior corona radiata R/L | 0.009/0.010 | 1085/661 |
| Superior corona radiata R/L | 0.009/0.010 | 852/561 |
| Posterior corona radiata R/L | 0.010/0.015 | 66/114 |
| External capsule R/L | 0.008/0.010 | 977/874 |
| Posterior limb of internal capsule R/L | 0.008/0.010 | 588/659 |
| Retrolenticular part of internal capsule R/L | 0.008/0.009 | 363/487 |
| Anterior limb of internal capsule R/L | 0.009/0.009 | 339/396 |
| Sagittal Stratum R/L | 0.008/0.010 | 382/230 |
| Posterior thalamic radiation R/L | 0.021/0.016 | 52/245 |
| Cerebral peduncle R/L | 0.008/0.009 | 321/217 |
| Corticospinal tract R/L | 0.008/0.009 | 130/141 |
| Medial lemniscus R/L | 0.008/0.008 | 118/136 |
| Pontine crossing tract | 0.008 | 281 |
| Fornix (cres) R/L | 0.008/0.011 | 264/154 |
| Uncinate fasciculus R/L | 0.008/0.010 | 44/33 |

**Figure C4.**
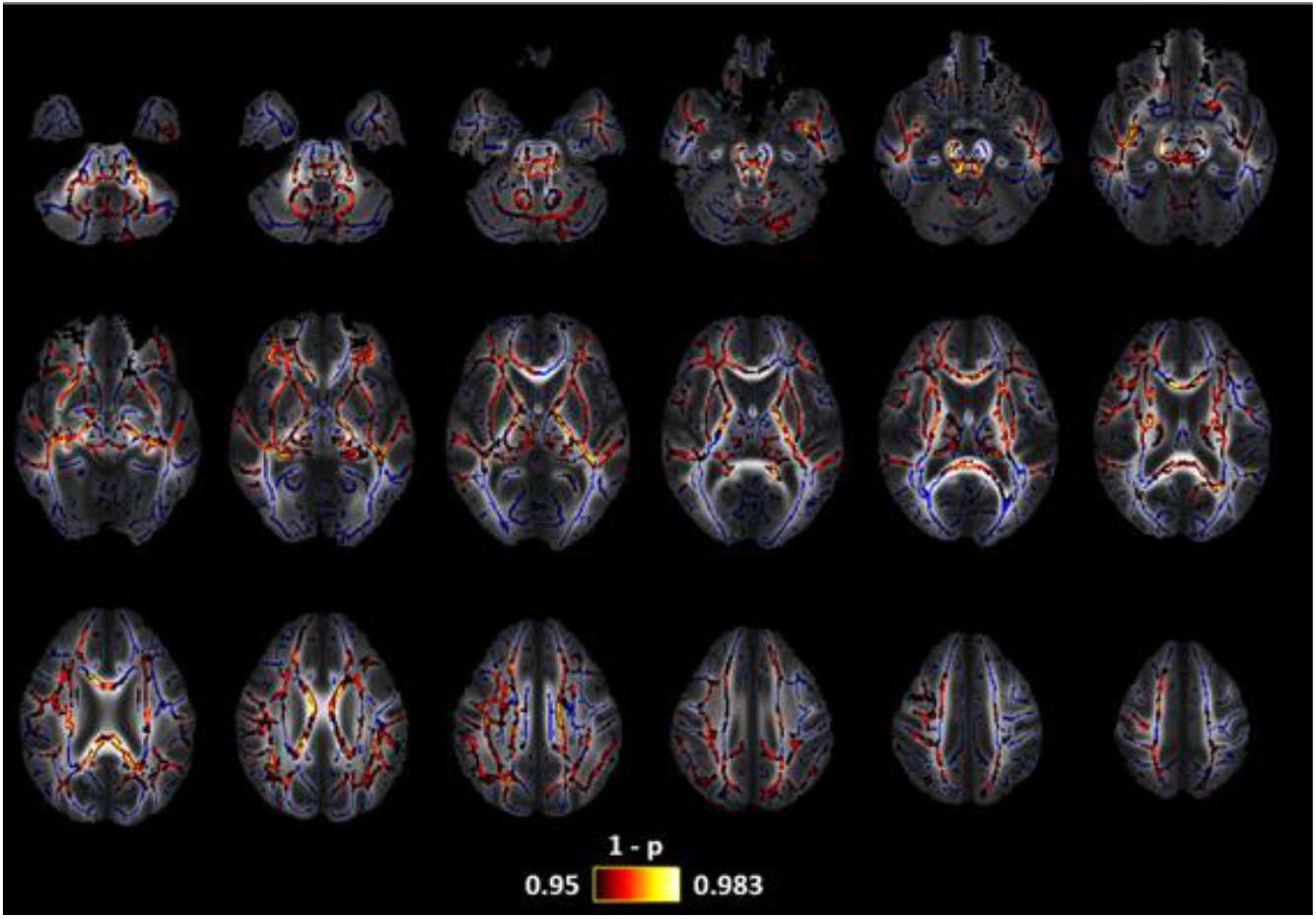
Return-to-axis (RTAP) changes in patients with EM in comparison with HC. Lower RTAP values were found in EM. The white matter skeleton is shown in blue and voxels with significant differences in red-yellow. The color bar shows the 1-p values (family-wise error corrected).

**Table C4.** White matter regions from the ICBM-DTI-81 White Matter Atlas for which significant decreased RTAP values were found in HC compared to EM.

| White Matter Region | Minimum *p*-Value (FWE-Corrected) | Volume (mm<sup>3</sup>) |
| --- | --- | --- |
| Middle cerebellar Peduncle | 0.027 | 1131 |
| Superior cerebellar peduncle R/L | 0.025/0.026 | 91/83 |
| Inferior cerebellar peduncle R/L | 0.035/0.037 | 119/37 |
| Superior fronto-occipital fasciculus R/L | 0.022/0.034 | 296/42 |
| Superior longitudinal fasciculus R/L | 0.022/0.032 | 654/296 |
| Genu of corpus callosum | 0.019 | 459 |
| Body of corpus callosum | 0.029 | 1343 |
| Splenium of corpus callosum | 0.041 | 1164 |
| Anterior corona radiata R/L | 0.016/0.036 | 1031/395 |
| Superior corona radiata R/L | 0.020/0.035 | 668/4430 |
| Posterior corona radiata R/L | 0.041/0.039 | 53/69 |
| External capsule R/L | 0.018/0.035 | 921/759 |
| Posterior limb of internal capsule R/L | 0.025/0.033 | 486/615 |
| Retrolenticular part of internal capsule R/L | 0.022/0.033 | 190/387 |
| Anterior limb of internal capsule R/L | 0.021/0.036 | 318/413 |
| Sagittal Stratum R/L | 0.022/0.035 | 298/136 |
| Cerebral peduncle R/L | 0.027/0.027 | 310/105 |
| Corticospinal tract R/L | 0.026/0.028 | 87/97 |
| Medial lemniscus R/L | 0.026/0.026 | 109/115 |
| Pontine crossing tract | 0.025 | 206 |
| Fornix (cres) R/L | 0.022/0.032 | 236/93 |
| Uncinate fasciculus R/L | 0.023/0.037 | 51/35 |

**Figure C5.**
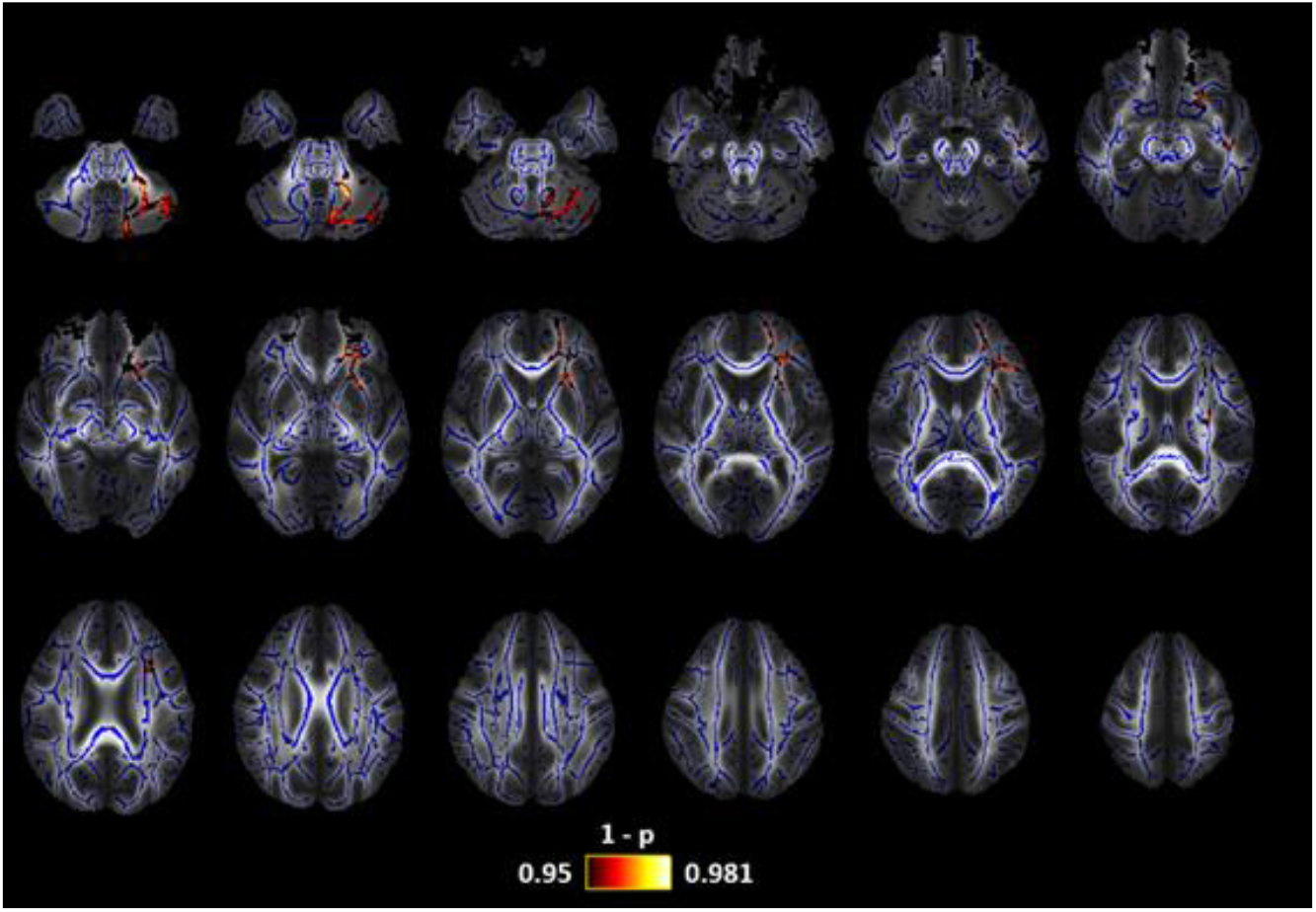
Return-to-plane (RTPP) changes in patients with EM in comparison with HC. Lower RTPP values were found in EM. The white matter skeleton is shown in blue and voxels with significant differences in red-yellow. The color bar shows the 1-p values (family-wise error corrected).

**Table C5.** White matter regions from the ICBM-DTI-81 White Matter Atlas for which significant decreased RTPP values were found in HC compared to EM.

| White Matter Region | Minimum $p$-Value (FWE-Corrected) | Volume (mm<sup>3</sup>) |
| --- | --- | --- |
| Middle cerebellar Peduncle | 0.019 | 521 |
| Anterior corona radiata L | 0.041 | 251 |
| External capsule L | 0.045 | 145 |
| Anterior limb of internal capsule R/L | 0.045 | 45 |

**Figure C6.**
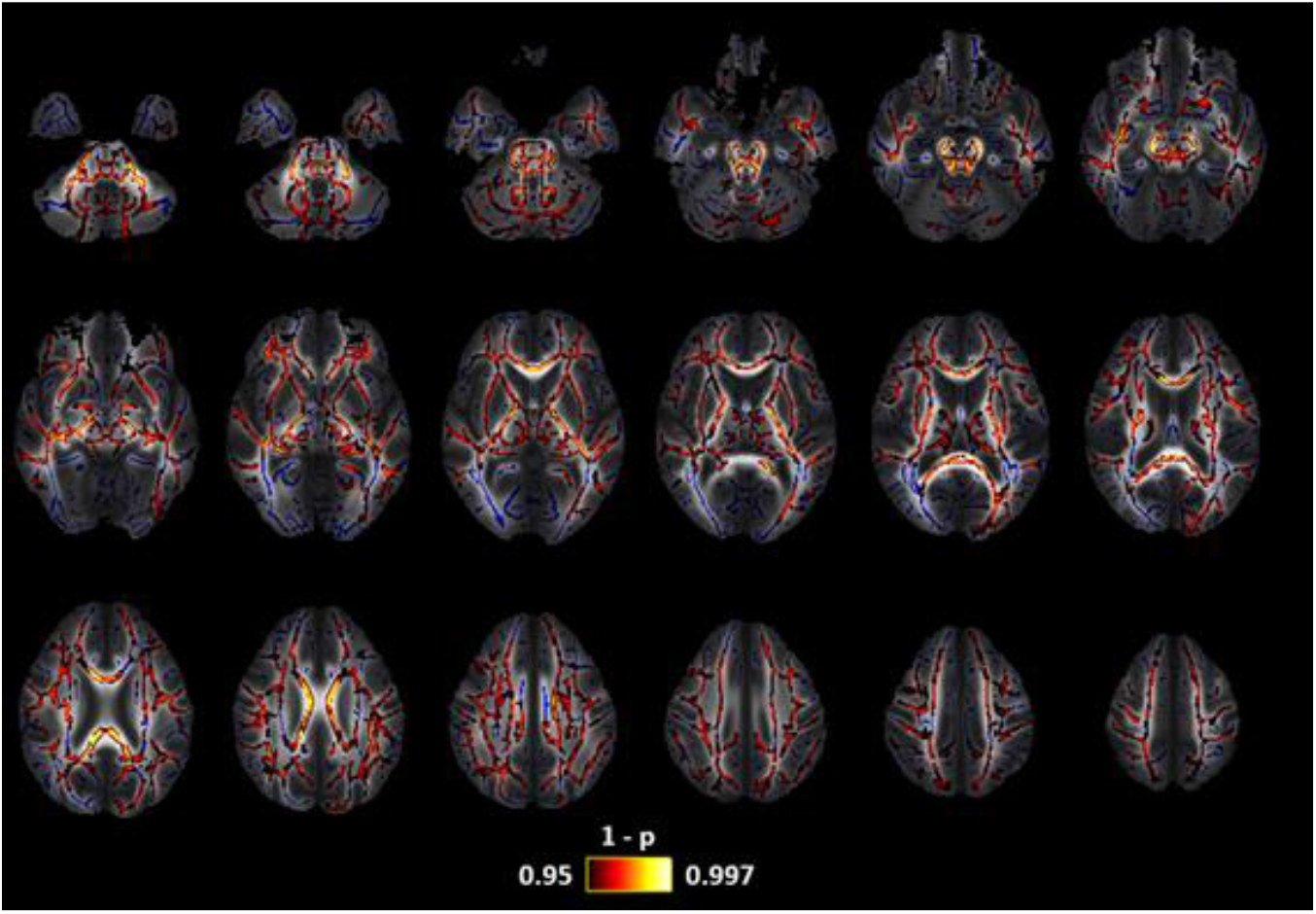
Q-Space Mean Square Displacement (qMSD) changes in patients with EM in comparison with HC. Lower qMSD values were found in EM. The white matter skeleton is shown in blue and voxels with significant differences in red-yellow. The color bar shows the 1-p values (family-wise error corrected).

**Table C6.** White matter regions from the ICBM-DTI-81 White Matter Atlas for which significant decreased qMSD values were found in HC compared to EM.

| White Matter Region | Minimum $p$-Value (FWE-Corrected) | Volume (mm<sup>3</sup>) |
| --- | --- | --- |
| Middle cerebellar Peduncle | 0.002 | 2082 |
| Superior cerebellar peduncle R/L | 0.002/0.002 | 186/148 |
| Inferior cerebellar peduncle R/L | 0.002/0.003 | 168/137 |
| Superior fronto-occipital fasciculus R/L | 0.003/0.004 | 47/44 |
| Superior longitudinal fasciculus R/L | 0.004/0.004 | 919/799 |
| Genu of corpus callosum | 0.004 | 1077 |
| Body of corpus callosum | 0.005 | 1964 |
| Splenium of corpus callosum | 0.004 | 1891 |
| Anterior corona radiata R/L | 0.004/0.004 | 1142/649 |
| Superior corona radiata R/L | 0.004/0.003 | 866/637 |
| Posterior corona radiata R/L | 0.005/0.004 | 108/195 |
| External capsule R/L | 0.002/0.004 | 1083/958 |
| Posterior limb of internal capsule R/L | 0.002/0.004 | 717/744 |
| Retrolenticular part of internal capsule R/L | 0.002/0.004 | 417/502 |
| Anterior limb of internal capsule R/L | 0.003/0.004 | 397/463 |
| Sagittal Stratum R/L | 0.002/0.004 | 397/226 |
| Posterior thalamic radiation R/L | 0.005/0.005 | 53/231 |
| Cerebral peduncle R/L | 0.002/0.003 | 487/400 |
| Corticospinal tract R/L | 0.002/0.003 | 170/209 |
| Medial lemniscus R/L | 0.002/0.002 | 159/154 |
| Pontine crossing tract | 0.002 | 316 |
| Fornix (cres) R/L | 0.002/0.004 | 297/187 |
| Cingulum (cyngulate gyrus) L | 0.008/0.008 | 86/97 |
| Uncinate fasciculus R/L | 0.002/0.004 | 47/39 |

**Figure C7.**
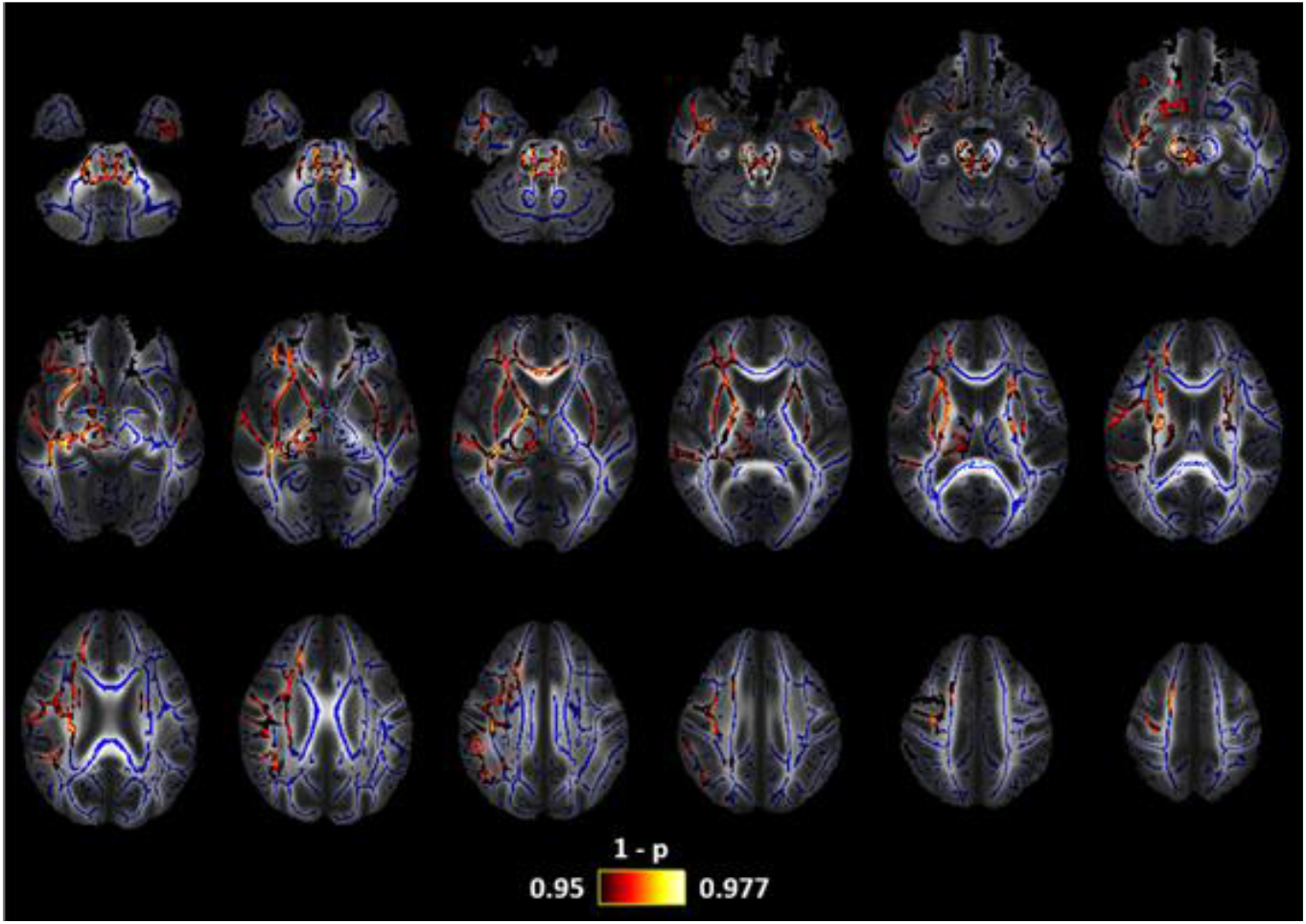
Propagator Anisotropy (APA) changes in patients with EM in comparison with HC. Lower APA values were found in EM. The white matter skeleton is shown in blue and voxels with significant differences in red-yellow. The color bar shows the 1-p values (family-wise error corrected).

**Table C7.**
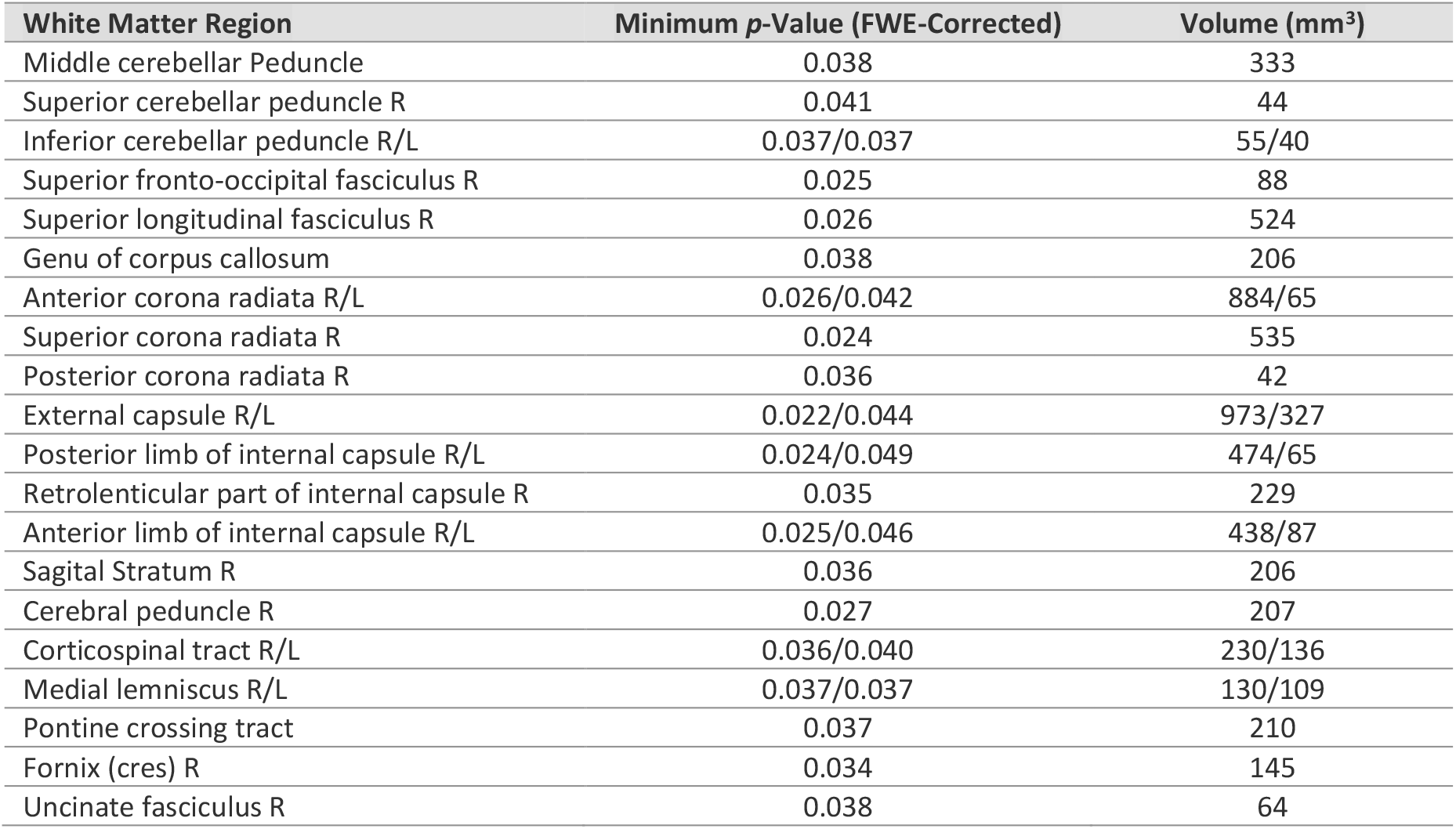
White matter regions from the ICBM-DTI-81 White Matter Atlas for which significant decreased APA values were found in HC compared to EM.

| White Matter Region | Minimum *p*-Value (FWE-Corrected) | Volume (mm<sup>3</sup>) |
| --- | --- | --- |
| Middle cerebellar Peduncle | 0.038 | 333 |
| Superior cerebellar peduncle R | 0.041 | 44 |
| Inferior cerebellar peduncle R/L | 0.037/0.037 | 55/40 |
| Superior fronto-occipital fasciculus R | 0.025 | 88 |
| Superior longitudinal fasciculus R | 0.026 | 524 |
| Genu of corpus callosum | 0.038 | 206 |
| Anterior corona radiata R/L | 0.026/0.042 | 884/65 |
| Superior corona radiata R | 0.024 | 535 |
| Posterior corona radiata R | 0.036 | 42 |
| External capsule R/L | 0.022/0.044 | 973/327 |
| Posterior limb of internal capsule R/L | 0.024/0.049 | 474/65 |
| Retrolenticular part of internal capsule R | 0.035 | 229 |
| Anterior limb of internal capsule R/L | 0.025/0.046 | 438/87 |
| Sagittal Stratum R | 0.036 | 206 |
| Cerebral peduncle R | 0.027 | 207 |
| Corticospinal tract R/L | 0.036/0.040 | 230/136 |
| Medial lemniscus R/L | 0.037/0.037 | 130/109 |
| Pontine crossing tract | 0.037 | 210 |
| Fornix (cres) R | 0.034 | 145 |
| Uncinate fasciculus R | 0.038 | 64 |

**Figure C8.**
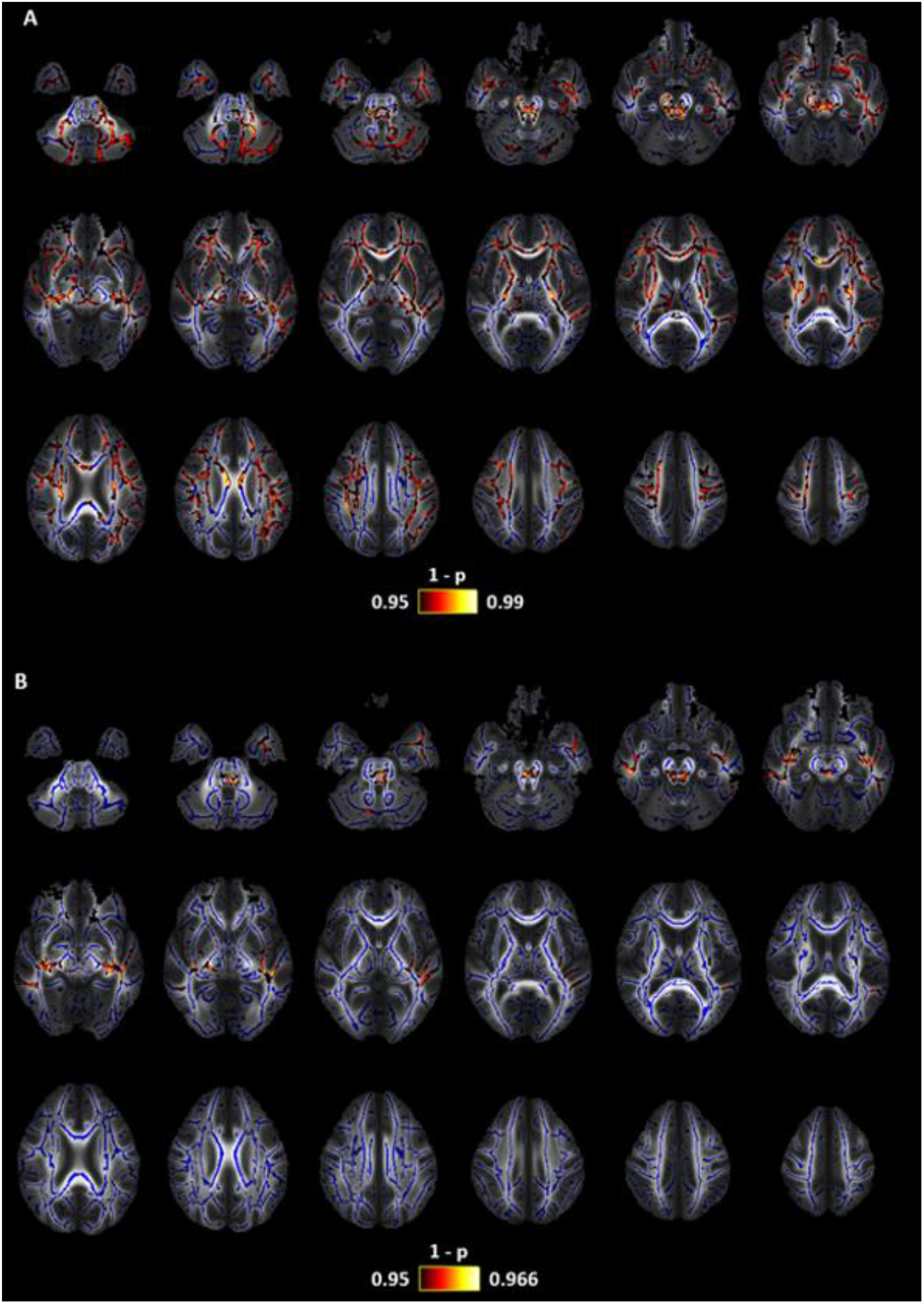
Moment 1/2 of the Ensemble Average Propagator (**ϒ**^1/2^) changes. A) Patients with CM in comparison with EM. Lower MUA values were found in EM. B) HC in comparison with EM patients. Lower **ϒ**^1/2^ values were found in EM. The white matter skeleton is shown in blue and voxels with significant differences in red-yellow. The color bar shows the 1-p values (family-wise error corrected).

**Table C8.** White matter regions from the ICBM-DTI-81 White Matter Atlas for which significant decreased **ϒ**^1/2^ values were found in CM compared to EM and HC compared to EM.

| HC compared to EM |  |  |
| --- | --- | --- |
| White Matter Region | Minimum $p$-Value (FWE-Corrected) | Volume (mm<sup>3</sup>) |
| External capsule R/L | 0.044/0.035 | 80/179 |
| Retrolenticular part of internal capsule L | 0.039 | 39 |
| Sagittal Stratum R/L | 0.041/0.036 | 113/118 |
| Medial lemniscus L | 0.043 | 44 |
| Pontine crossing tract | 0.042 | 107 |
| Fornix (cres) R/L | 0.045/0.036 | 35/74 |
| CM compared to EM |  |  |
| Middle cerebellar Peduncle | 0.009 | 1324 |
| Superior cerebellar peduncle R/L | 0.029/0.029 | 54/97 |
| Inferior cerebellar peduncle L | 0.013 | 46 |
| Superior fronto-occipital fasciculus R | 0.024 | 37 |
| Superior longitudinal fasciculus R/L | 0.023/0.023 | 694/734 |
| Genu of corpus callosum | 0.020 | 488 |
| Body of corpus callosum | 0.023 | 548 |
| Splenium of corpus callosum | 0.048 | 66 |
| Anterior corona radiata R/L | 0.038/0.020 | 488/630 |
| Superior corona radiata R/L | 0.023/0.023 | 750/330 |
| Posterior corona radiata R/L | 0.041/0.033 | 55/56 |
| External capsule R/L | 0.023/0.021 | 478/615 |
| Posterior limb of internal capsule R/L | 0.023/0.026 | 443/271 |
| Retrolenticular part of internal capsule L | 0.028 | 205 |
| Anterior limb of internal capsule R/L | 0.027/0.021 | 297/188 |
| Sagittal Stratum R/L | 0.034/0.025 | 152/227 |
| Cerebral peduncle R | 0.032 | 238 |
| Corticospinal tract R/L | 0.032 | 36 |
| Medial lemniscus R/L | 0.033 | 72 |
| Pontine crossing tract | 0.029 | 87 |
| Fornix (cres) R/L | 0.035/0.029 | 103/91 |
| Fornix (column and body) | 0.041 | 41 |

**Figure C9.**
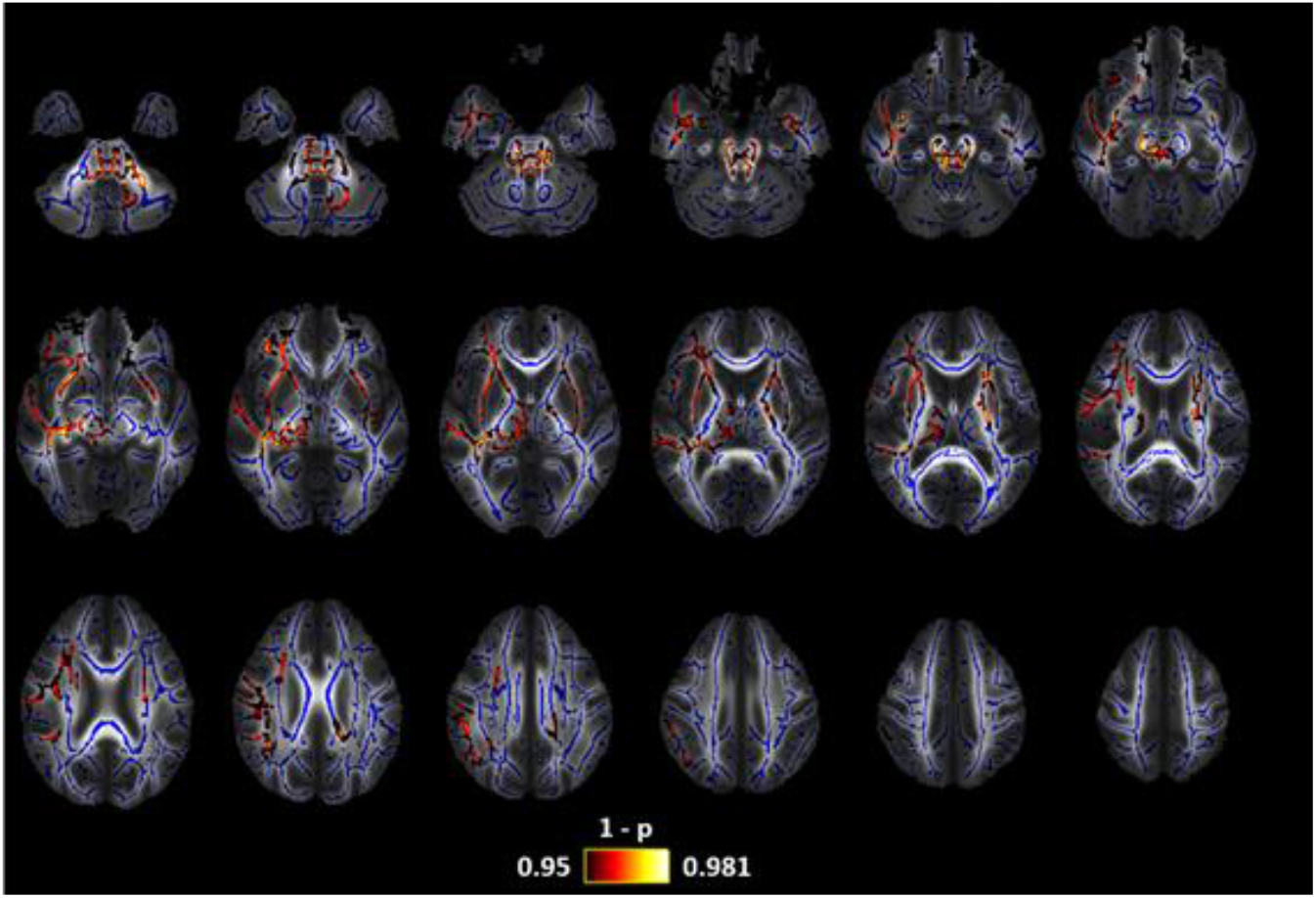
Diffusion Anisotropy (DiA) changes in patients with EM in comparison with HC. Lower DiA values were found in EM. The white matter skeleton is shown in blue and voxels with significant differences in red-yellow. The color bar shows the 1-p values (family-wise error corrected).

**Table C9.** White matter regions from the ICBM-DTI-81 White Matter Atlas for which significant decreased DiA values were found in EM compared to HC.

| White Matter Region | Minimum $p$-Value (FWE-Corrected) | Volume (mm<sup>3</sup>) |
| --- | --- | --- |
| Middle cerebellar Peduncle | 0.020 | 546 |
| Superior cerebellar peduncle R/L | 0.020/0.021 | 94/78 |
| Inferior cerebellar peduncle R/L | 0.019/0.020 | 72/43 |
| Superior fronto-occipital fasciculus R | 0.020 | 64 |
| Superior longitudinal fasciculus R | 0.019 | 368 |
| Body of corpus callosum | 0.044 | 38 |
| Splenium of corpus callosum | 0.044 | 59 |
| Anterior corona radiata R/L | 0.019/0.042 | 695/53 |
| Superior corona radiata R/L | 0.020/0.045 | 248/95 |
| External capsule R/L | 0.018/0.042 | 956/382 |
| Posterior limb of internal capsule L | 0.042 | 206 |
| Retrolenticular part of internal capsule R | 0.019 | 320 |
| Anterior limb of internal capsule R/L | 0.019/0.042 | 242/137 |
| Sagittal Stratum R | 0.019 | 265 |
| Cerebral peduncle R/L | 0.019/0.024 | 243/74 |
| Corticospinal tract R/L | 0.019/0.019 | 224/160 |
| Medial lemniscus R/L | 0.018/0.019 | 148/132 |
| Pontine crossing tract | 0.020 | 154 |
| Fornix (cres) R | 0.019 | 193 |
| Uncinate fasciculus R | 0.022 | 62 |

## Appendix D. Resampling of diffusion measures

As commented in section 2.4.2 and 3.2, this experiment was also carried out for p < 0.01. Figure D1 shows the results.

**Figure D1.**
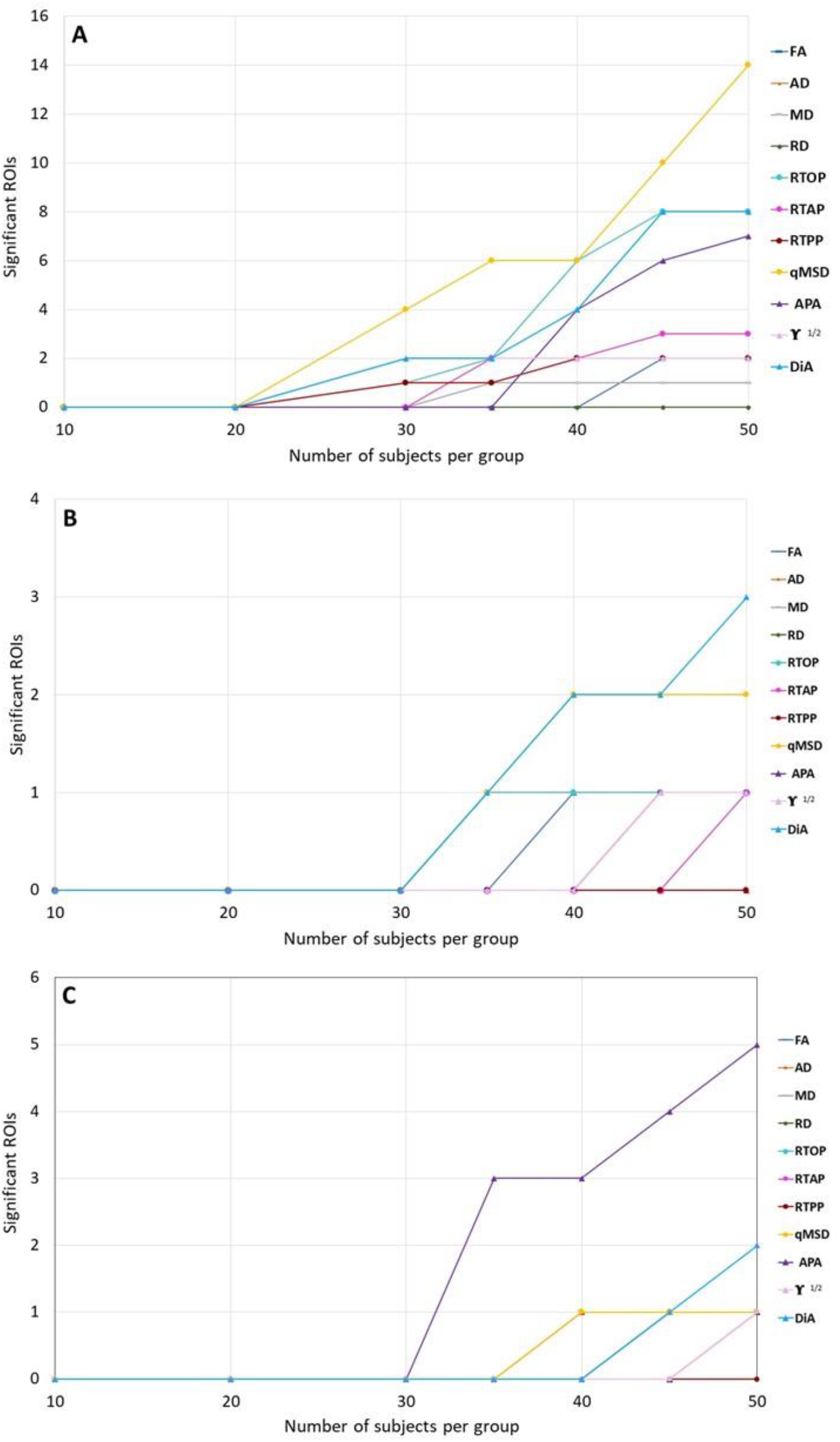
Significant ROIs by the resampling of diffusion measures reducing the number of subjects per group for p < 0.01. A) Episodic Migraine (EM) vs. Healthy Controls (HC). B) Chronic Migraine (CM) vs. HC. C) CM vs. EM. DTI (triangles) and AMURA (circles) measures are shown in the graph.

